# NALCN/Cch1 channelosome subunits originated in early eukaryotes and are fully conserved in animals, fungi, and apusomonads

**DOI:** 10.1101/2025.04.04.647097

**Authors:** Adriano Senatore, Tatiana D. Mayorova, Luis Yanez-Guerra, Wassim Elkhatib, Brian Bejoy, Philippe Lory, Arnaud Monteil

## Abstract

The sodium leak channel NALCN, a key regulator of neuronal excitability, associates with three ancillary subunits that are critical for its function: an extracellular subunit called FAM155, and two cytoplasmic subunits called UNC79 and UNC80. Interestingly, NALCN and FAM155 have orthologous phylogenetic relationships with the fungal calcium channel Cch1 and its extracellular subunit Mid1, however, UNC79 and UNC80 have not been reported outside of animals. In this study, we leveraged expanded gene sequence data available for eukaryotes to re-examine the evolutionary origins of NALCN and Cch1 channel subunits. Our analysis corroborates the direct phylogenetic relationship between NALCN and Cch1 and identifies a larger clade of related channels in additional eukaryotic taxa. We also identify homologues of FAM155/Mid1 in Cryptista algae, and UNC79 and UNC80 homologues in numerous non-metazoan eukaryotes including basidiomycete and mucoromycete fungi, and the microbial eukaryotic taxa Apusomonadida, Malawimonadida, and Discoba. Furthermore, we find that most major animal lineages, except ctenophores, possess a full complement of NALCN subunits. Comparing structural predictions with the solved structure of the human NALCN complex supports orthologous relationships between metazoan and non-metazoan FAM155/Mid1, UNC79, and UNC80 homologues. Together, our analyses reveal unexpected diversity and ancient eukaryotic origins of NALCN/Cch1 channelosome subunits and raise interesting questions about the functional nature of this conserved channel complex within a broad, eukaryotic context.

## Introduction

The sodium leak channel NALCN represents a fourth major branch of four-domain cation channel in animals, the others being low-voltage activated calcium channels (*i.e.*, Ca_V_3 channels), high voltage activated calcium channels (Ca_V_1 and Ca_V_2), and voltage-gated sodium channels (Na_V_)^1,2^. Studies in several animals, primarily in nematode worms, fruit flies, and mice have revealed critical functions for NALCN in processes including sleep, circadian rhythm, breathing, nociception, pain, locomotion, and parturition^3–5^. At the cellular level, NALCN contributes depolarizing leak sodium currents that help set the resting membrane potential of neurons and other excitable cells, and its activity can be modulated to exert changes in cellular excitability^3^.

Both *de novo* dominant and inherited recessive pathogenic variants of NALCN are described in severe pathological conditions in humans characterized by a wide range of symptoms, and NALCN is implicated in many other diseases including psychiatric disorders and cancer^3^. In vivo and in vitro, NALCN does not operate on its own but as part of a large multi-protein complex herein referred to as the NALCN channelosome. Specifically, NALCN associates with an extracellular subunit called FAM155, and two large cytoplasmic subunits, UNC79 and UNC80^6–8^. These additional ancillary proteins are necessary for the functional expression of NALCN, its trafficking to the plasma membrane, its cellular localization, and indeed stability of the entire channelosome complex (reviewed in^3–5^).

Interestingly, previous analyses revealed that NALCN and FAM155 have phylogenetic orthologues in fungi, where they are referred to as Cch1 and Mid1, respectively^1,2,9^. In fungi, Cch1 and Mid1 form a highly regulated Ca^2+^ permeable channel that restores calcium levels in response to various stimuli including endoplasmic reticulum stress and pheromone signaling, a function that contributes to virulence of several pathogenic fungi including *Cryptococcus neoformans*, *Candida albicans*, and *Aspergillus funigatus*^10–14^. Instead, UNC79 and UNC80, which form a heterodimer in the cytoplasm, have not been reported outside of animals. In this study, we took advantage of the expanded genome and transcriptome sequence data that is currently available to re-interrogate the presence of NALCN subunits in animals, fungi, and other eukaryotes. Our results reveal an unexpected diversity of ancillary NALCN/Cch1 channelosome subunits in eukaryotes, including complete conservation of a tetrameric complex between animals and fungi.

## Results

### Phylogeny of the NALCN and Cch1 subunits

Previous phylogenetic analyses demonstrated homology between NALCN and Cch1 channels^1,2^. We sought to expand on this previous work by searching for homologues in a collection of 185 high-quality proteomes from a range of species spanning eukaryotes, utilizing a custom Hidden Markov Model strategy (*i.e.*, with HMMer3) which is more sensitive than BLAST for identifying homologues^15^. Thus, we generated an HMM profile trained on a set of manually selected NALCN and Cch1 channel protein sequences and used this to identify a total of 3,476 non-redundant protein sequences from our proteome collection. These were then filtered to remove ones with less than 18 predicted transmembrane helices, to exclude single and tandem domain pore-loop channels like voltage-gated potassium and two-pore channels, respectively. The resulting 1,476 candidate four domain channel sequences were then analyzed with the all-against-all sequence similarity clustering program CLANS. This revealed a main cluster of 1,454 sequences each with a minimum of three connections with other proteins, comprised of large central group which includes metazoan Ca_V_1/Ca_V_2, Ca_V_3, and Na_V_ channels, together with a diverse set of non-metazoan channels (Figure 1). NALCN channels from animals, along with fungal Cch1 channels, formed a separate and highly connected subcluster, and several additional subclusters were evident for sets of channels from the stramenopile, alveolate, and rhizaria (SAR) supergroup, cryptista, and chloroplastida. Of note, since our HMM model trained on NALCN /Cch1 sequences was also able to detect other classes of distant four domain channels, including Ca_V_ and Na_V_, it seems unlikely that our search strategy missed detection of NALCN/Cch1 sequences within the utilized proteomes.

**Figure 1.**
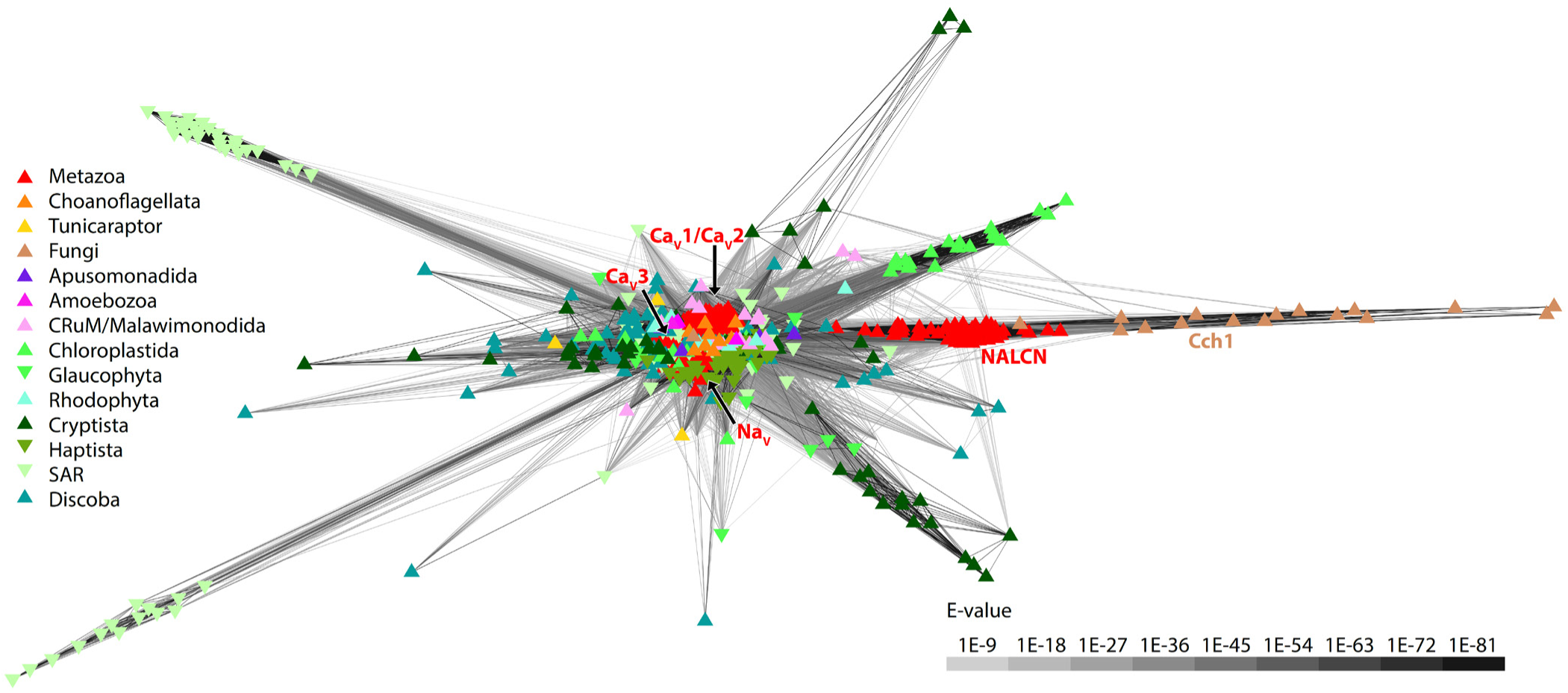
PAM30 all vs. all cluster map of eukaryotic pore-loop channels bearing more than 18 transmembrane helices identified with a hidden Markov model trained with metazoan NALCN and fungal Cch1 proteins sequences. Edges correspond to BLAST comparison expect-values (E-values) colored according to the provided legend on the lower right of the map. Individual Ion channel sequences are depicted by symbols which are colored according to the taxonomic groups indicated by the legend on the left of the plot. Metazoan NALCN, voltage-gated sodium (Na_V_) and calcium (Ca_V_) channels, as well as fungal Cch1 channels, are labeled.

To explore the relationships of these HMM-identified channels, we generated a maximum likelihood phylogeny inferred from a trimmed protein sequence alignment. The resulting tree confirms the direct phylogenetic relationship between NALCN and Cch1 channels^1,2^, and identifies a strongly supported sister clade relationship of NALCN/Cch1 with channels from species within Apusomonadida and Crumalia/CRuM (*i.e.*, collodictyonids, rigifilids, and mantamonadids) (Figure 2A). Together, these associate with a larger set of channels from Discoba, Malawimonadida, and Cryptista, in a clade with variable node support, and a much broader strongly supported clade, we refer to as clade A, with representation amongst eukaryotes. Metazoan Ca_V_1 and Ca_V_2 channels, Ca_V_3 channels, and Na_V_ channels fall within three distant and separate clades from NALCN/Cch1 channels and indeed all clade A channels (*i.e.*, clades B, C, and D respectively) (Figure 2A), with each clade also containing other sets of channels from microbial eukaryotic taxa including Choanoflagellata, Chloroplastida, Cryptista, and Apusomonadida.

**Figure 2.**
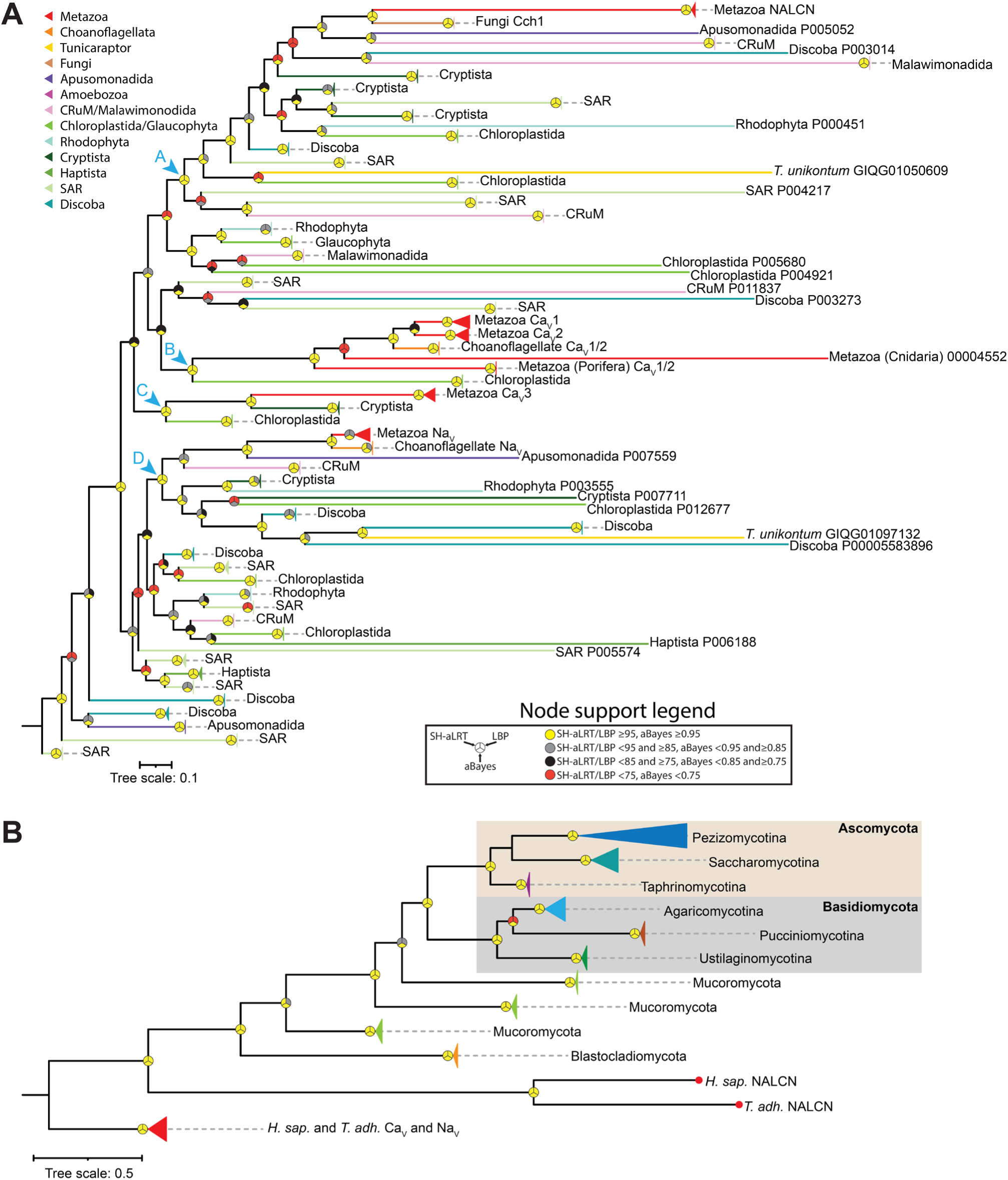
Phylogenetic analysis of eukaryotic pore-loop channels related to NALCN and Cch1. **A)** Maximum likelihood phylogenetic tree of HMM-identified pore-loop channel protein sequences from a curated set of eukaryotic proteomes. Branches are colored according to taxonomic groupings as indicated by the legend. The chevrons with alphabetic labels (cyan colored) denote strongly supported nodes that separate NALCN/Cch1 (clade A), Ca_V_1/Ca_V_2 (clade B), Ca_V_3 (clade C), and Na_V_ (clade D) and associated eukaryotic channels from each other. **B)** Maximum likelihood phylogenetic tree of select metazoan pore-loop channels and Cch1 homologues identified from an expanded set of fungal proteomes from the FungiDB database. Leaves on the tree are colored according to major taxonomic groupings within fungi. For both trees, node support values for three separate analyses, SH-aLRT, LBP, and aBayes, are depicted by circular symbols with colors reflecting ranges of values as indicated in the node support legend.

Next, we used our NALCN/Cch1 Hidden Markov Model to search though an expanded set of 256 proteomes from the FungiDB genomic database, which includes species from all the major fungal lineages: Ascomycota, Basidiomycota, Blastocladiomycota, Chytridiomycota, and Mucoromycota^16^. This enabled us to identify Cch1 homologues in all lineages except Chytridiomycota (filtered to possess at least 18 predicted transmembrane helices). A phylogenetic tree of these protein sequences, rooted on human and *Trichoplax adhaerens* Ca_V_ and Na_V_ channels, reveals near congruency with the expected species phylogeny^17^, except for a switched branch position of Cch1 homologues from Pucciniomycotina and Ustilagomycotina within the larger taxonomic group of Basidiomycota (Figure 2B).

We also conducted a more detailed phylogenetic analysis of NALCN in animals, using a reciprocal BLAST search approach to identify homologues in gene data from representative species spanning most major bilaterian and non-bilaterian phyla (Figure S1, inset). Except for ctenophores (comb jellies), we were able to identify NALCN homologues in all animal taxa. A phylogenetic tree of these protein sequences reveals most examined species possess only single gene copies of NALCN, with independent duplications apparent for the known *Caenorhabditis elegans* paralogues NCA-1 and NCA-2, the platyhelminth *Macrostomum lignano*, and the great barrier reef sponge *Amphimedon queenslandica* (Figure S1).

### FAM155 homologues are found in animals, fungi, and unicellular cryptist algae

Like Cch1, previous phylogenetic analyses established a phylogenetic link between FAM155 in animals and Mid1 in fungi^9^. Using the same strategy that we used for NALCN and Cch1 channels, we generated an HMM model trained on selected FAM155 and Mid1 sequences and used it to search through the set of 185 eukaryotic proteomes. This identified a total of 44 protein sequences, mostly from animals and fungi as expected. However, as is evident in a sequence similarity cluster map, eight of these sequences were from algae-like species from the group Cryptista, with closer connections to fungal Mid1 sequences compared to FAM155 sequences from metazoans (Figure 3A). Of note, our HMM search failed to identify a previously identified FAM155/Mid1 homologue from the apusomonad species *Thecamonas trahens*^9^ (NCBI accession number XP_013754286.1), despite the sequence being present in the proteome we used in our analysis. We attribute this to the generally divergent nature of FAM155/Mid1 proteins^9^, and the atypical length of this protein of 2909 amino acids, compared, for example, to only 458 for human FAM155A and 623 for *C. neoformans* (fungal) Mid1 (see discussion). A maximum likelihood phylogeny based on the identified sequences reveals strong support for nodes separating metazoan, fungal, and cryptist FAM155/Mid1 homologues, but weak or absent support for most nodes within each taxon (Figure 3B). We used this same HHM model to search through our expanded set of fungal proteomes, identifying homologues in all lineages except Chytridiomycota, like the Cch1 subunit. Hence, it seems all fungal lineages that possess Cch1 also possess Mid1.

**Figure 3.**
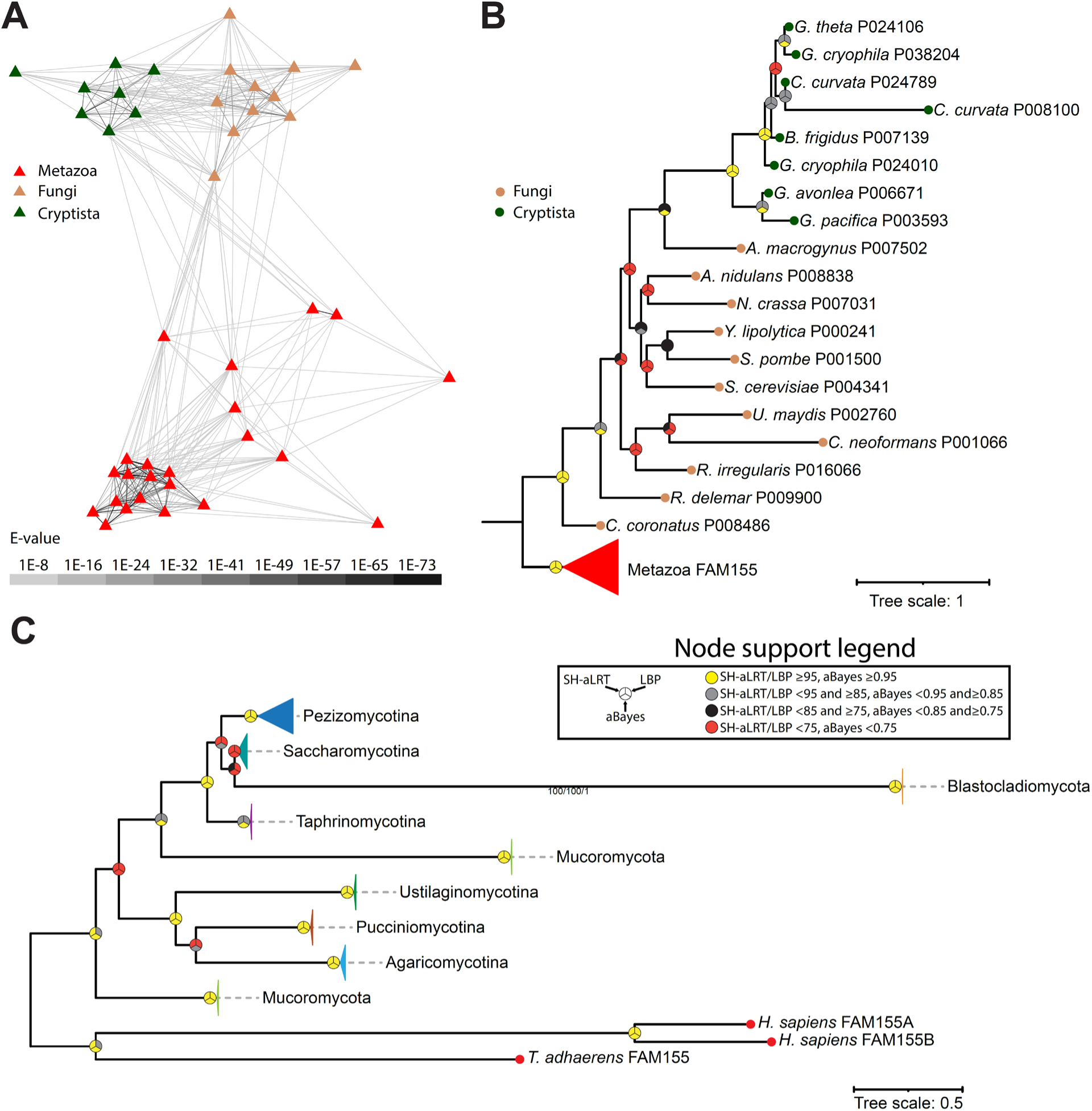
FAM155/Mid1 homologues are found in metazoa, fungi, and cryptista. **A)** BLOSSUM45 all vs. all cluster map of HMM-identified FAM155/Mid1 proteins sequences from eukaryotes. Edges correspond to BLAST comparison e-values colored according to the legend. **B)** Maximum likelihood tree of HMM-identified FAM155/Mid1 homologues in eukaryotes. **C)** Maximum likelihood tree of HMM-identified FAM155/Mid1 homologues in an expanded set of fungal proteomes. Leaves on the tree are colored according to major taxonomic groupings within fungi. For both trees, node support values for three separate analyses, SH-aLRT, LBP, and aBayes, are depicted by circular symbols with colors reflecting ranges of values as indicated in the node support legend.

Like NALCN, searching for FAM155 sequences in animals via reciprocal BLAST allowed us to identify homologues in all major taxa except ctenophores. A phylogenetic tree based on these sequences suggests independent duplications occurred in vertebrates (*i.e.*, FAM155A and FAM155B), the rotifer *Rotaria socialis*, and the platyhelminth *M. lignano* that also duplicated NALCN (Figure S2).

Our identification of FAM155/Mid1 homologues in cryptists prompted us to explore their homology with metazoan FAM155 and fungal Mid1 proteins more deeply, along with the previously identified homologue from *T. trahens*. Using AlphaFold3^18^, we predicted dimeric complexes of NALCN-FAM155 proteins from the early-diverging animal *T. adhaerens*, and the Cch1-Mid1 homologues from the basidiomycete fungus *C. neoformans*, in which these have been shown to be functionally and genetically integrated^11,19,20^. We also included in these analysis Mid1/Fam155 homologues from two cryptist species, *Cryptomonas curvata* and *Geminigera cryophila*, and the apusomonad *T. trahens*. Putative NALCN/Cch1 homologues for these species were selected from our phylogenetic analysis of four-domain channels, specifically, from the clades of cryptist and apusomonad channels most phylogenetically proximal to NALCN and Cch1 (Figure 2A)^21^.

Although such structural predictions must be interpreted with caution, we reasoned that AlphaFold could be used to explore whether complexing, in a manner consistent with the known structures of human NALCN and FAM155, is at least possible for these various proteins.

Unfortunately, AlphaFold failed to predict the dimeric structure of the *T. trahens* NALCN/Cch1 and FAM155/Mid1 proteins, producing a low predicted template modeling (pTM) score of 0.45 (all scores and accession numbers for our structural predictions are provided in Supplementary Table 2). Nonetheless, all other examined dimers produced acceptable scores above 0.5, all containing a set of α1 to α3 helices positioned atop the channel (Figure 4A and B), which in the solved human complex mediates critical contacts with the NALCN subunit ^22–24^. We also predicted the structure of the human NALCN-FAM155A complex and structurally aligned it with its corresponding solved structure^7^, recealing highly overlapping structures with a root mean square alignment deviation score of 0.784 angstroms (Figure S3). Outside of the α1 to α3 helices, there is marked variation in the predicted structures of FAM155/Mid1 proteins with a set of β sheets in the *T. adhaerens* subunit, an extended globular arrangement in the *C. neoformans* subunit, and various flanking alpha helices in the *G. cryophila* subunit. Notably, the *T. adhaerens* FAM155 subunit is also predicted to possess a hydrophobic alpha helix that runs alongside the NALCN subunit (Figure 4A).

**Figure 4.**
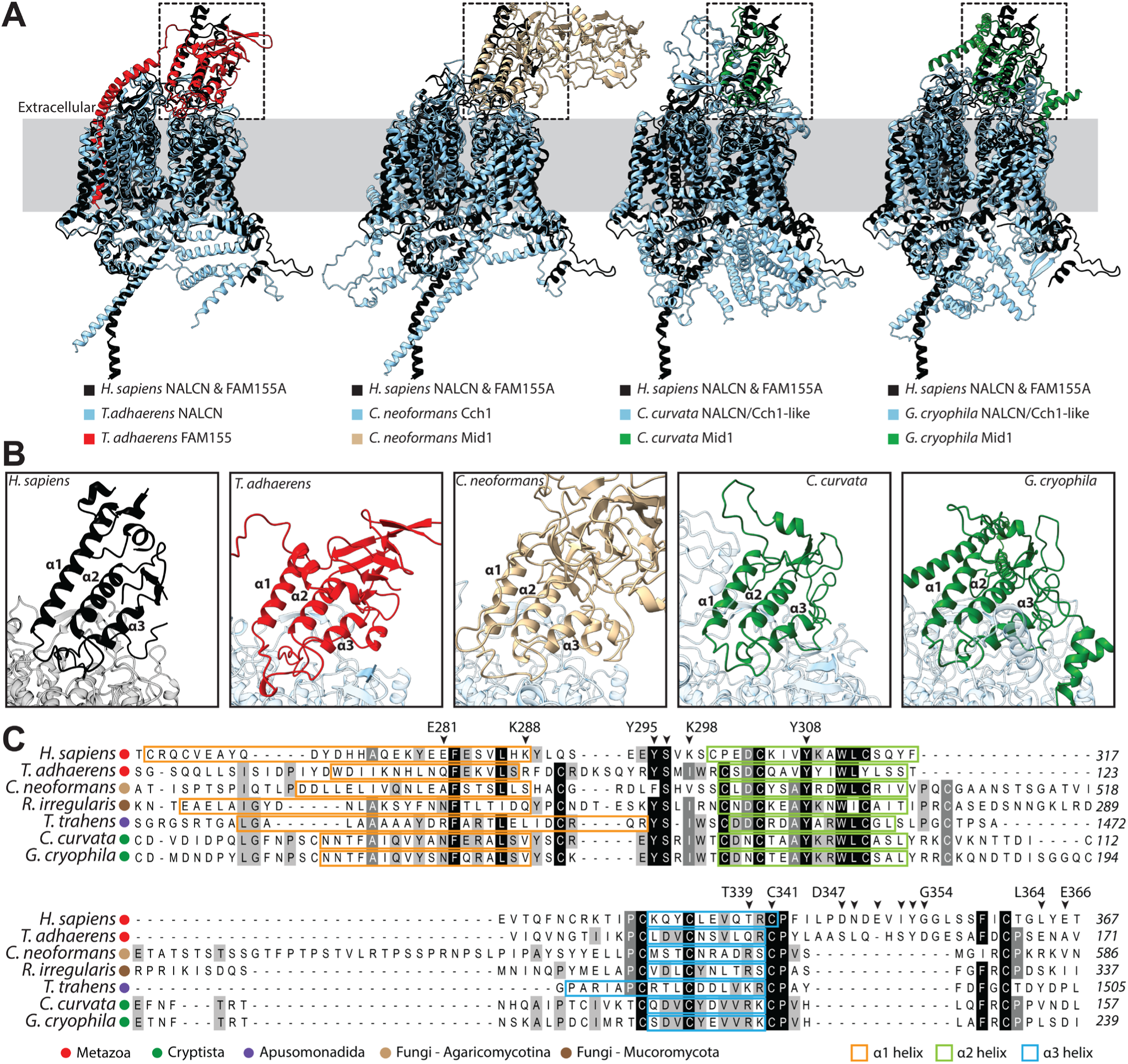
Structural and sequence analysis of FAM155/Mid1 homologues. **A)** Structural superimposition of the solved human NALCN-FAM155A cryo-EM structure (from PDB number 7XS4) and predicted NALCN-FAM155 or CCH1-Mid1 dimer structures from the early-diverging animal *Trichoplax adhaerens*, the pathogenic fungus *Cryptococcus neoformans*, and the cryptista (algal) species *Cryptomonas curvata* and *Geminigera cryophila*. **B)** Close-up view of the FAM155/Mid1 NALCN/CCH1 subunit interfaces (corresponding to the dashed black boxes in panel A), revealing similar positioning of predicted α1 to α3 helices that in the human complex form critical contacts with the NALCN subunit. **C)** Partial protein alignment of selected FAM155/Mid1 homologues from animals, fungi, apusomonads, and cryptists, revealing strong sequence conservation within α1 to α3 helices. The chevrons above the alignment denote amino acids in the human FAM155A shown to make important contacts with NALCN in cryo-EM structures. Numbers to the right of the alignment denote amino acid positions. The boxes are used to label amino acids predicted to form helical structures by AlphaFold.

A protein alignment of human FAM155A with homologues from *T. adhaerens*, the fungi *C. neoformans* and *R. irregularis*, the apusomonad *T. trahens*, and the cryptists *C. curvata* and *G. cryophila* reveals conserved sites in α1 to α3 helices, within otherwise highly divergent protein sequences (Figure 4C). Among the three helices, α1 is the most divergent both in terms of sequence and length (*i.e.*, 16 to 38 amino acids). The α2 and α3 helices exhibit more conservation, most possessing highly conserved cysteine residues that serve to stabilize the tertiary arrangement of the α1 to α3 helices in resolved structures^22–24^. Several amino acids within and around these alpha helices form critical contacts with NALCN in mammals, but our alignment shows that most of these are not conserved. The exceptions are a tyrosine-serine motif in the α1-α2 linker, a tyrosine residue in α2, and a cysteine in α3. Thus, it appears considerable changes have occurred in molecular determinants that mediate the interaction between NALCN/Cch1 and FAM155/Mid1 subunits. Also notable is that the region located downstream of α3 (*i.e.*, the post α3 loop), which is believed to be essential for interactions with NALCN in mammals, is poorly conserved in non-vertebrate sequences beyond a ubiquitously conserved cysteine-proline motif at the very start of this linker (Figure 4C).

### UNC79 and UNC80 homologues are found within several distant eukaryotic lineages

To date, unlike NALCN/CCH1 and FAM155/Mid1, bona fide UNC79 and UNC80 homologues have not been identified in neither fungi nor any other non-metazoan eukaryotes. Nonetheless, given the expanded set of gene sequences available for fungi and other eukaryotes, we sought to re-examine the presence of these two genes outside of animals. A BLAST search through the NCBI non-redundant database identified putative UNC80 homologues from several fungal species including *Rhizopus delemar* (Mucoromycota; NCBI accession number KAG1050648.1). A reciprocal BLAST of this sequence against metazoan NCBI sequences identified UNC80 homologues as top hits, and global alignment of this protein with the human UNC80 sequence revealed a low percent identity of 22.3% and percent similarity of 37.1% (File S1). With a stringent expect value cutoff of 1E-30, we used the *R. delemar* sequence to extract additional homologues from the FungiDB database and combined these with a set of metazoan UNC80 homologues to train a Hidden Markov Model for searching through our set of eukaryotic proteomes. This led to the identification of 139 UNC80 homologues, most from metazoans, but also from fungi, Rotosphaerida, Apusomonadida, Malawimonadida, CRuMs, and Discoba (Figure 5A). A phylogeny of these proteins reveals strong support for the separation of animal UNC80 sequences from those of other eukaryotes, and respective monophyletic relationships among the five fungal sequences and the two from Discoba.

**Figure 5.**
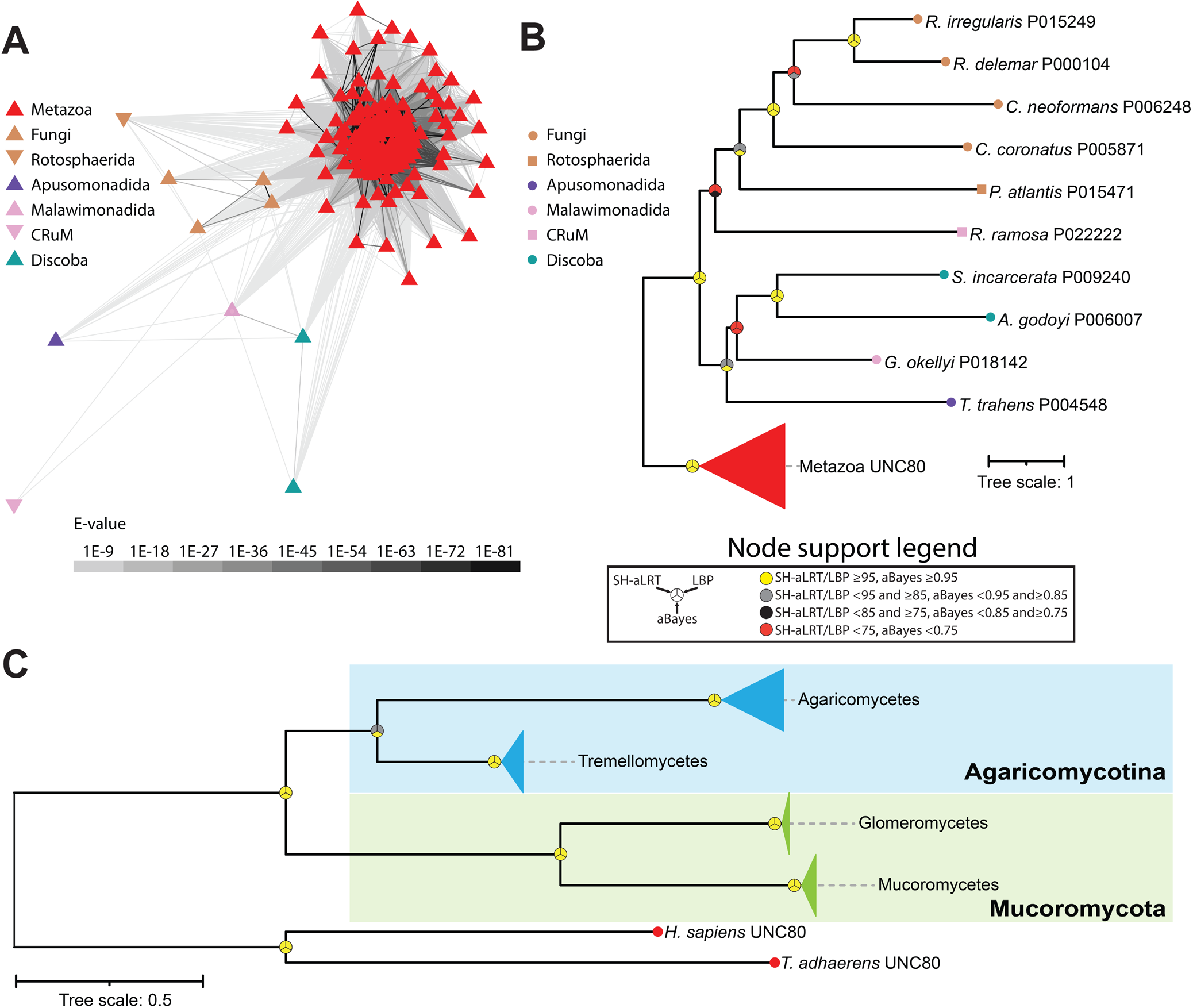
Identification of UNC80 homologues in fungi and other eukaryotes. **A)** BLOSSUM62 all vs. all cluster map of HMM-identified UNC80 proteins sequences from eukaryotes. Edges correspond to BLAST comparison e-values colored according to the legend. **B)** Maximum likelihood tree of HMM-identified UNC80 homologues in eukaryotes. **C)** Maximum likelihood tree of HMM-identified UNC80 homologues in an expanded set of fungal proteomes. Leaves on the tree are colored according to major taxonomic groupings within fungi. For both trees, node support values for three separate analyses, SH-aLRT, LBP, and aBayes, are depicted by circular symbols with colors reflecting ranges of values as indicated in the node support legend.

To better examine the presence of UNC80 in fungi, we generated two separate HMM models, one of combined metazoan and fungal sequences, and another with just fungal sequences, and used these to search through the set of FungiDB proteomes. While both models identified a common set of 42 candidate homologues, the fungi-only model identified two additional sequences. Thus, we selected this latter set of 44 sequences for phylogenetic analysis. A maximum likelihood tree, rooted on the human and *T. adhaerens* UNC80 homologues, reveals narrower representation among fungal species compared to Cch1 and Mid1, restricted to agaricomycetes and tremellomycetes within the Basidiomycota, Agaricomycotina, and glomeromycetes and mucoromycetes within Mucoromycota. Focusing on animals using the reciprocal BLAST strategy, we were able to identify UNC80 homologues in all examined species except ctenophores, thus similar to NALCN and FAM155, with an apparent duplication of this subunit in *R. socialis* (Figure S4).

Unlike UNC80, a BLAST search for UNC79 homologues outside of animals failed to produce any hits in the NCBI non-redundant database. However, while conducting these analyses, a preprint article was released documenting the identification of putative UNC79 and UNC80 homologues in the basidiomycete *C. neoformans*. Through large scale genotype-phenotype clustering, this study found a phenotypic link between the previously characterized Cch1 and Mid1 genes, and two previously uncharacterized and large proteins bearing predicted armadillo repeats, a feature of UNC79 and UNC80^25^. This study also demonstrated a biochemical interaction between these two proteins, and with both Cch1 and Mid1, altogether suggesting these form a complex with Cch1 and are homologous to the animal UNC79 and UNC80 subunits of NALCN^25^. However, phylogenetic evidence supporting this homology was not generated.

Using the putative *C. neoformans* UNC79 homologue, we identified several additional fungal sequences via a BLAST search of the FungiDB database (using an expect value cutoff of 1E-30) and combined a selection of these with metazoan UNC79 sequences for training an HMM model and used this combined model to search for homologues in our eukaryotic databases. This identified a total of 140 sequences, which interestingly, were similar to UNC80 in terms of species composition, most coming from animals and fungi, but also species from Apusomonadida, Malawimonadida, and Discoba (Figure 6A and B). Our inability to detect fungal UNC79 sequences via BLAST, using animal UNC79 sequences as queries, indicates strong sequence divergence. A global alignment of the *C. neoformans* and human UNC79 proteins yields only 23.1% sequence identity and 37.8% sequence similarity, distributed broadly along the length of the alignment without any extended stretches of matching sequence (Supplementary File 1). This is different from our alignment of the *R. delemar* and human UNC80 (Supplementary File 1), and likely accounts for the inability of BLAST to seed and extend a high scoring pair based on metazoan UNC79 query sequences. Considering this, we decided to generate an HMM model of UNC79 based exclusively on the manually selected fungal sequences from our combined model and used this to search for homologues in the FungiDB data sets. This yielded a total 39 hits, which when analyzed phylogenetically produced a tree very similar to the fungal UNC80 phylogeny with respect to species composition and topology (Figures 5C and 6C). Evident from this analysis is that, like UNC80, UNC79 homologues are restricted to agaricomycetes and tremellomycetes within the Agaricomycotina, and glomeromycetes and mucoromycetes within the Mucoromycota. Together, these analyses indicate that select clades of fungi within Agaricomycotina and Mucoromycota possess a complete set of NALCN channelosome subunits: NALCN/Cch1, FAM155, UNC79, and UNC80. Lastly, reciprocal BLAST and phylogenetic analysis of UNC79 homologues, focused strictly within animals, identified sequences in all examined species except ctenophores (Figure S5), altogether indicating that all animals, except ctenophores, possess a complete set NALCN subunits.

**Figure 6.**
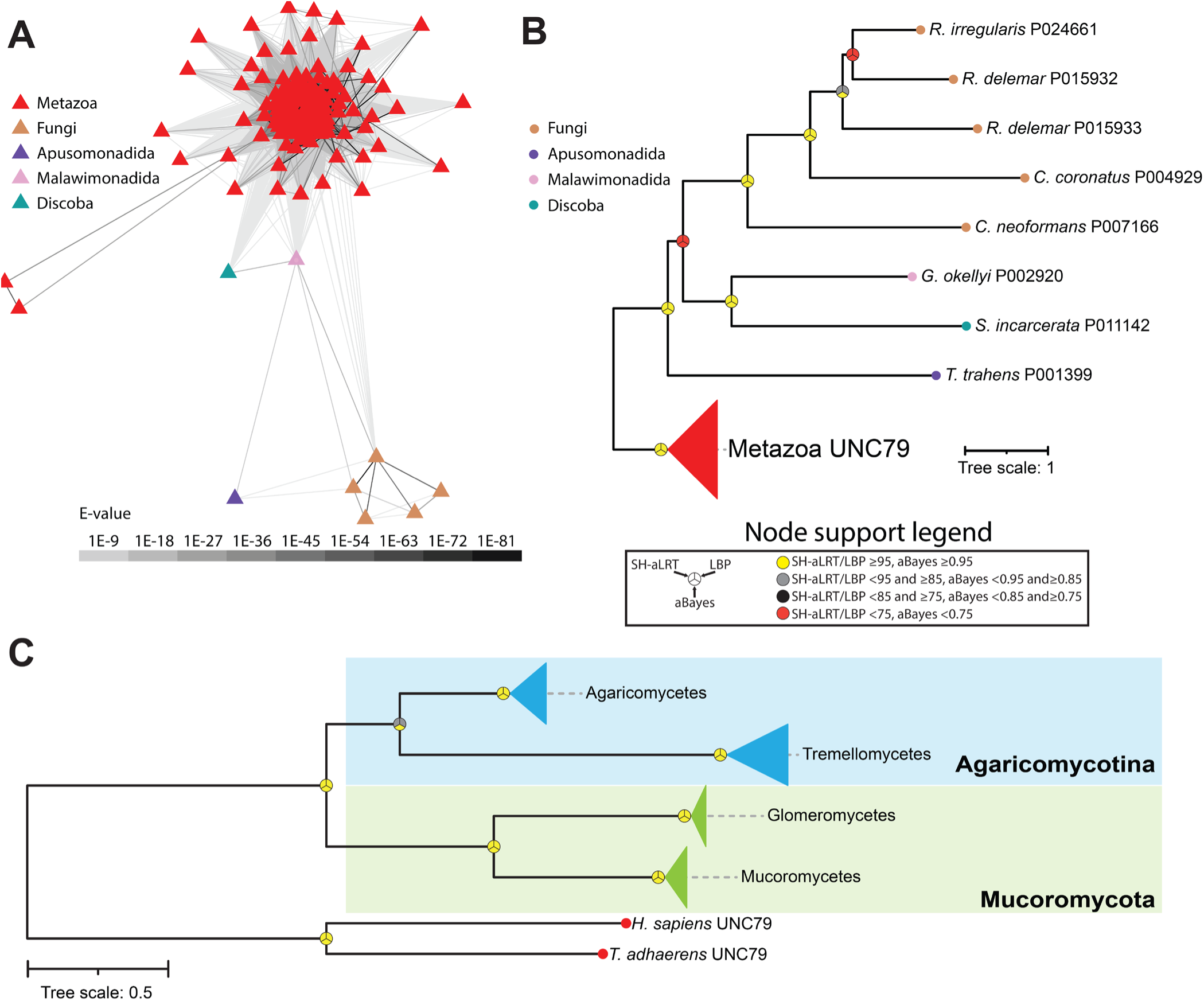
Identification of UNC79 homologues in fungi and other eukaryotes. **A)** BLOSSUM62 all vs. all cluster map of HMM-identified UNC79 proteins sequences from eukaryotes. Edges correspond to BLAST comparison e-values colored according to the legend. **B)** Maximum likelihood tree of HMM-identified UNC79 homologues in eukaryotes. **C)** Maximum likelihood tree of HMM-identified UNC79 homologues in an expanded set of fungal proteomes. Leaves on the tree are colored according to major taxonomic groupings within fungi. For both trees, node support values for three separate analyses, SH-aLRT, LBP, and aBayes, are depicted by circular symbols with colors reflecting ranges of values as indicated in the node support legend.

Clearly, both the identified non-metazoan UNC79 and UNC80 homologues are highly divergent in their protein sequence relative to their metazoan counterparts. In cryo-EM structures, the human subunits both form longitudinal, S-shaped proteins each composed of more than 30 armadillo repeats, which together form a sub-dimer in an inverted N-to C-terminal supercoiled orientation^6–8^ (Figure 7A). We sought to predict whether such configurations are possible for the identified non-metazoan homologues using AlphaFold3. Specifically, we predicted the dimeric structures of homologues from the species *T. adhaerens*, *C. neoformans* (fungi, Agaricomycotina), *R. irregularis* (fungi, Mucoromycota), *Thecamonas trahens* (Apusomonadida), *Gefionella okellyi* (Malawimonadida), and *Stygiella incarcerata* (Discoba). All proteins were predicted to form armadillo repeat proteins, with dimers forming inverted N-to C-terminal arrangements. However, only the homologues from *T. adhaerens*, *C. neoformans*, *R. irregularis*, *T. trahens*, *G. okellyi* had linear supercoiled structures that could be structurally aligned with the solved structures of human UNC79 and UNC80 (Figure 7B to F). Instead, the predicted dimer from *S. incarcerata* lacked supercoiling (Figure 7G). Thus, the structural predictions are consistent with the ability of these proteins to form dimeric complexes like the known structure of human UNC79 and UNC80, further support the homology of these non-metazoan proteins to metazoan NALCN subunits, and are consistent with the recent biochemical evidence that the UNC79 and UNC80 homologues from *C. neoformans* form a functional and physical complex with Cch1 and Mid1^25^.

**Figure 7.**
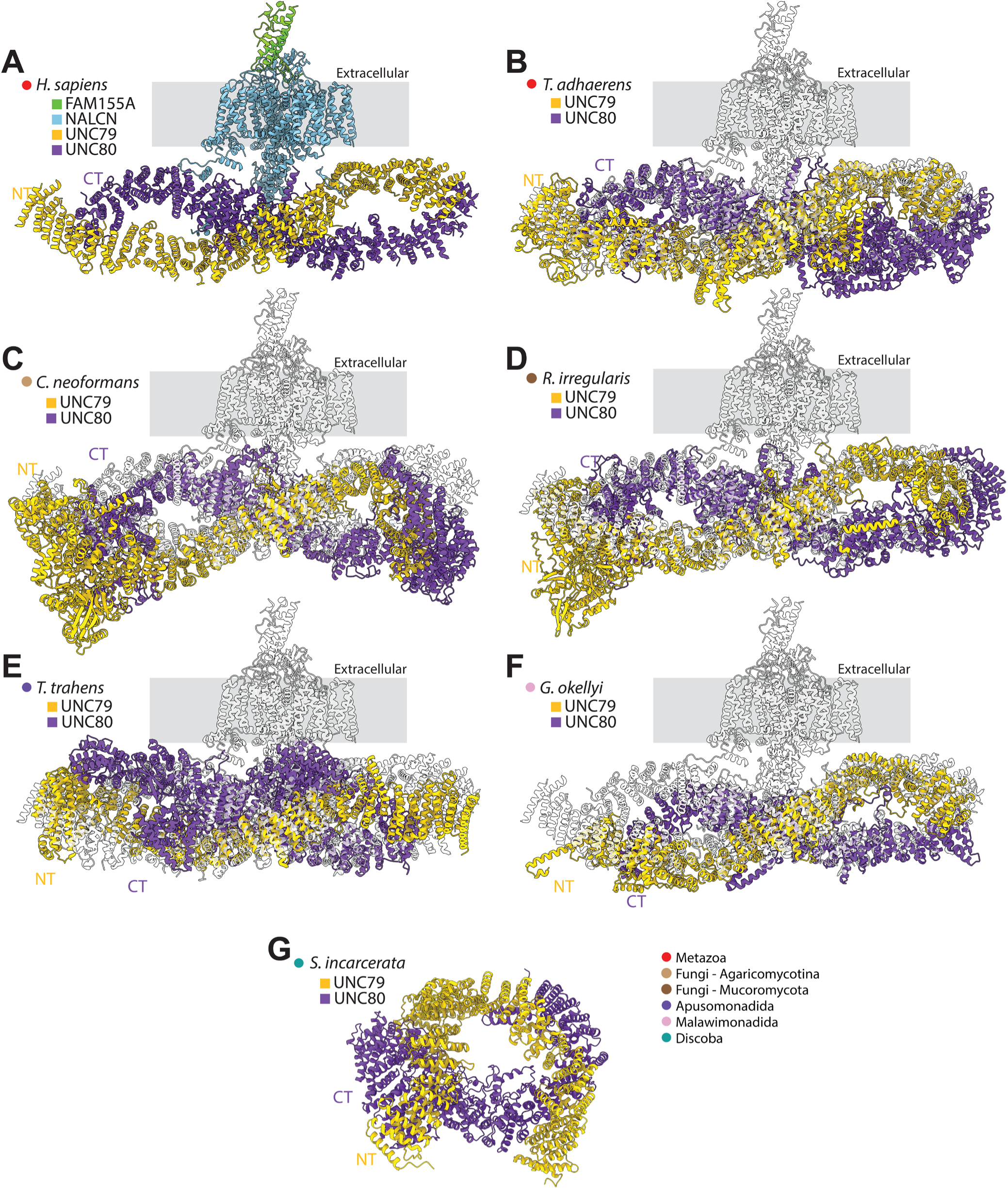
Structural analysis of UNC79 and UNC80 homologues from animals and other eukaryotes. **A)** Cryo-EM structure of the human NALCN channel complexed with the extracellular subunit FAM155A and the cytoplasmic subunits UNC79 and UC80 (PDB accession number 7XS4). **B)** Structural superimposition of human NALCN complex (in semi-transparent white) with the predicted structure of the UNC79/UNC80 dimer from the early-diverging metazoan *Trichoplax adhaerens*. **C)** Structural superimposition of human NALCN complex with the predicted structure of the UNC79/UNC80 dimer from the fungal species *Cryptococcus neoformans*. **D)** Structural superimposition of human NALCN complex with the predicted structure of the UNC79/UNC80 dimer from the fungal species *Rhizophagus irregularis*. **E)** Structural superimposition of human NALCN complex with the predicted structure of the UNC79/UNC80 dimer from the apusomonad *Thecamonas trahens*. **F)** Structural superimposition of human NALCN complex with the predicted structure of the UNC79/UNC80 dimer from the malawimonad *Gefionella okellyi*. **G)** Predicted structure of the UNC79/UNC80 dimer from the Discoba species *Stygiella incarcerata*. The legend in the lower right indicates relevant taxonomic groupings.

## Discussion

Our phylogenetic analysis of four domain channels identified using a custom HMM profile trained on NALCN and Cch1 protein sequences revealed that NALCN, Ca_V_1/Ca_V_2, Ca_V_3, and Na_V_ channels fall into four separate and strongly supported clades (*i.e.*, clades A to E), each comprised of additional non-metazoan channels from a broad range of eukaryotes (Figure 2A). In clade A, which included NALCN and Cch1, there were also channels from Apusomonadida, Malawimonadida, and Crumalia/CRuMs, which formed phylogenetic relationships with NALCN/Cch1 roughly consistent with the species phylogeny^21^ (Figure 8). Clade A also contained more divergent channels from eukaryotes within the Diaphoretikes (*i.e.*, Chloroplastida, Glaucophyta, Rhodophyta, Cryptista, SAR, and Discoba), indicating that NALCN and Cch1 belong to a large and ancient clade of four domain channels, conserved between Amorphea and Diaphoretikes, that is distinct from Ca_V_ and Na_V_ channels.

**Figure 8.**
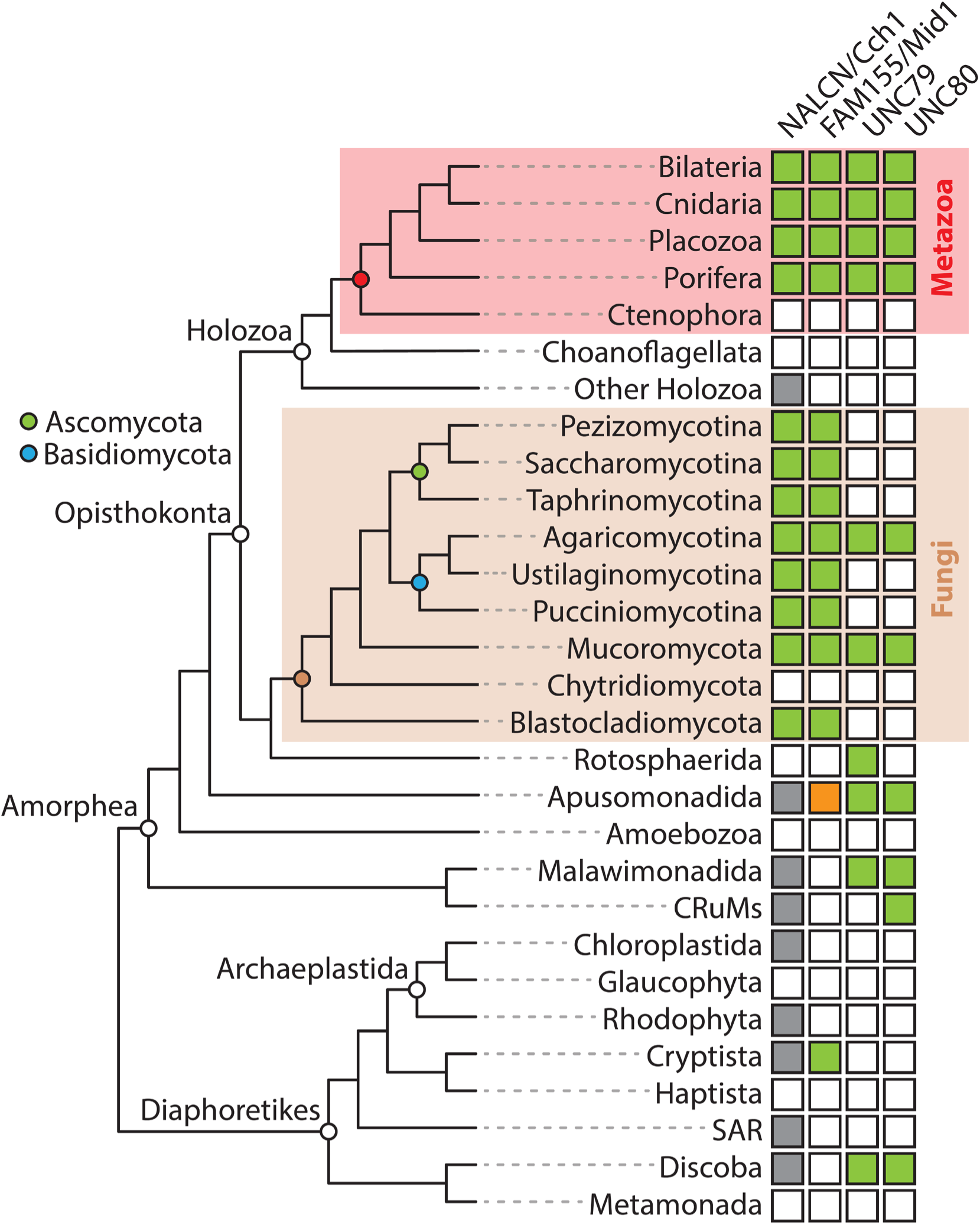
Summary of identified NALCN/CCH1 channelosome subunits in eukaryotes. Presence of homologues identified in this study are indicated by green colored boxes, with taxonomic groupings according recent phylogenomic studies^17,27,54,55^. Boxes colored in gray indicate the presence of NALCN/Cch1-related channels within clade A of our phylogenetic analysis shown in Figure 2A. The box colored in orange represents a FAM155/Mid1 homologue previously identified for the apusomonad *Thecamonas trahens*^9^.

Previous studies demonstrated orthologous phylogenetic relationships between metazoan NALCN and Mid1 proteins, and fungal Cch1 and Mid1 proteins, respectively^1,9^. Here, we extend these observations by demonstrating that two separate lineages of fungi, Agarocomycotina within the larger clade Basidiomycota, and Mucoromycota, also possess UNC79 and UNC80 homologues (Figure 8). We also found homologues of UNC79 and UNC80 more broadly in eukaryotes, within several non-opisthokont lineages within Amorphea, and Discoba within Diaphoretikes (Figure 8). Structural predictions of sets of these proteins from various representative species corroborated their homology to metazoan UNC79 and UNC80, most arranged as inverted N-to C-terminal supercoiled quaternary structures with each subunit made of repeating armadillo repeats (Figure 7), similar to the solved structures of human UNC79 and UNC80 in complex with NALCN and FAM155^6–8^ (Figure 8). That these are true homologues of metazoan UNC79 and UNC80 is supported by recent biochemical experiments revealing that Cch1, Mid1, UNC79, and UNC80 from the fungal species *C. neoformans* (Agaricomycota) form a physical complex in vitro, and large-scale mutation analysis indicating the four proteins form a functional phenotypic module in vivo^25^.

Outside of animals and fungi, the least prevalent subunit we found in our analysis was FAM155/Mid1 (Figure 8). However, we did identify bona fide homologues from Cryptista, with conserved α1 to α3 helices that form a core structural scaffold important for interactions with the NALCN subunit in solved structures of the mammalian channel complex^22–24^ (Figure 4). Worth noting is that FAM155/Mid1 proteins are relatively short in sequence and highly divergent outside of the α1 to α3 helices, and as such, our HMM search might have failed to identify some homologues. Indeed, this was the case for the FAM155/Mid1 homologue from the apusomonad *Thecamonas trahens* that was identified in a previous study^9^. Perhaps, our HMM model failed to identify this homologue not only because of sequence divergence, but also because of its unique extended length of 2909 amino acids, compared to the much shorter sequences for FAM155/Mid1 proteins we identified for animals, fungi, and cryptists (Figure 4C and Supplementary Files 10 and 11). Likewise, armadillo repeat proteins like UNC79 and UNC80 tend to be highly divergent, beyond retaining conserved helical tertiary structures^26^. Thus, it may be that sequences were missed, such that future analyses with improved taxon sampling combined with novel approaches might uncover additional FAM155/Mid1, UNC79, and UNC80 homologues in other eukaryotic lineages.

However, despite possible omissions, our combined analyses point to an ancient origin of the complete NALCN/Cch1 channelosome complex, with conservation of all four subunits in at least three eukaryotic lineages: metazoans, fungi, and apusomonads, all within Amorphea (Figure 8). In animals, for which our search was more extensive and directed, we found homologues of all NALCN channelosome subunits in all examined taxa except ctenophores (comb jellies), proposed to be the most early diverging^27,28^. Similarly in fungi, our deep search found Cch1 homologues in most phyla, all of which were also found to possess Mid1. However, as noted, only species from Mucoromycota and Agaricomycotina within Basidiomycota were found to possess UNC79 and UNC80 (Figure 8). Examining the presence/absence of genes strictly within opisthokonta, and assuming the absences are real, it seems all NALCN/Cch1 subunits were lost in the intervening lineages between animals and fungi, outside of a single channel found for the holozoan *Tunicaraptor unikontum*. Within fungi, UNC79 and UNC80 appear to have been independently lost in Ascomycota, Ustilagomycota, Pucciniomycota, and Blastocladiomycota, while Chytridiomycota lost the entire Cch1 channelosome complex (Figure 8).

Interestingly, the complete co-occurrence of all four subunits within animals, and of Cch1/Mid1 within fungi, suggests they have obligate functional relationship within each other, altogether consistent with the various genotype-phenotype studies that have been done in nematode worms, fruit flies, an mice (reviewed in^3–5^), and several species of fungi^10,20^. In contrast, and as noted, UNC79 and UNC80 are less prevalent in fungi, suggesting Cch1/Mid1 can function independently from UNC79/UNC80, permitting the loss of the latter subunits in select fungal lineages. Nonetheless, the recent evidence that *C. neoformans* Cch1, Mid1, UNC79, and UNC80 form a physical and functional complex^25^ suggests that fungal species possessing all four subunits have channels with similar obligate tetrameric relationships as observed in animals^3^.

Our analysis raises some interesting questions about conserved/divergent functions of these proteins alone or as a complex in these highly divergent eukaryotic contexts. In animals, the NALCN channelosome plays a conserved role in regulating the resting membrane potential of neurons and other excitable cells by conducting constitutive sodium currents that counter the hyperpolarizing potassium leak currents^3–5^. Emerging evidence indicates that NALCN is also subject to modulation hence serving as a ‘dial’ for regulating membrane potential and cellular excitation^3^. However, considering differences in key amino acids within the pore that define cation selectivity, it is unclear if this function is conserved in all animals. NALCN distinguishes itself from Ca_V_ and Na_V_ channels by having undergone extensive sequence divergence in the pore selectivity filter, a ring of four amino acids that serves to discriminate between permeating Na^+^, K^+^, and Ca^2+^ ions in all pore-loop channels. In general, the presence of glutamate and/or aspartate residues in the selectivity filter is associated with Ca^2+^ preference^29,30^, with Ca^2+^-selective channels like Ca_V_1 and Ca_V_2 bearing four glutamates (*i.e.*, selectivity filters of EEEE), or Ca_V_3 channels bearing two glutamates and two aspartates (EEDD). True sodium-selective Na_V_ channels on the other hand are thought to have evolved from Ca^2+^ selective channels with selectivity filter motifs of aspartate, glutamate, glutamate, alanine (DEEA), via mutation of the third glutamate to a positively charged lysine in a clade of bilaterian channels called Na_V_1 (*i.e.*, selectivity filters of DEKA), and independently in a set of cnidarian Na_V_ channels by a glutamate/lysine conversion in the second position (DKEA)^31^. The presence of a positively charged lysine in the selectivity filter is thought to disrupt the high affinity binding of Ca^2+^ ions in the pore, thus shifting the preference towards permeating Na^+^ ions^32^. Fungal Cch1 channels possess glutamate-rich selectivity filters like those of Ca_V_ channels^1^, consistent with their characterization as Ca^2+^ permeating channels^10–14^. Instead, NALCN channels have hypervariable selectivity filters that are subject to modification by alternative splicing. That is, while vertebrates, fruit flies, and nematode worms possess invariable NALCN selectivity filters of EEKE, consistent with sodium permeation, the most early diverging animals possess motifs of EEEE, consistent with calcium permeation^33^. In hemichordates, echinoderms, molluscs, and annelids, alternative splicing can produce NALCN channels with EEEE or EKEE selectivity filters, while in select arthropods, alternative splicing can produce channels with EEEE or EEKE selectivity filters^33^. To date, the ion selectivity properties of invertebrate NALCN channels with Ca_V_-like selectivity filters of EEEE have not been characterized, though based on the conserved significance of the selectivity filter for defining ion selectivity among Ca_V_, Na_V_, NALCN, and indeed all pore-loop channels, it seems likely that ancestral NALCN channels were more Ca^2+^ selective. Furthermore, like Na_V_ channels, the emergence of preferred Na^+^ permeation, via glutamate to lysine conversions in positions 2 or 3 of the selectivity filter, appears to have been a secondary adaptation that evolved independently several times in select animals^33^. Taken together, these features of the NALCN pore raise questions about the role the channel plays in different animals, either as a sodium leak channel, or potentially as a calcium permeating channel as observed in yeast.

Lastly, our identification of UNC79 and UC80 homologues in fungi and other eukaryotes raises interesting questions about the conserved functions of these two large proteins in the cytoplasm. We know little about their molecular functions in animals^3^, and even less in fungi or other eukaryotes^25^, and it may be that work in fungi or other non-metazoans might uncover some conserved functions that are relevant for NALCN biology in animals.

## Methods

### Sequence identification

We trained custom Hidden Markov Models (HMM) to search for homologues of NALCN, FAM155/Mid1, UNC79, and UNC80 within a set of high-quality eukaryotic proteomes spanning Amorphea (*e.g.*, animals, choanoflagellates, and fungi; 88 proteomes) and Diaphoretickes (plants and stramenopiles; 96 species), and separately, the complete set of fungal proteomes available in the integrated and functional genomic database FungiDB (release 68^16^). For the eukaryotic proteomes, details about sources and BUSCO^34^ quality metrics are provided in Supplementary Table 1. Collections of metazoan and fungal sequences were used to train the HMM models, manually extracted from available sequence data of various animal and fungal species in the NCBI non-redundant^35^ and FungiDB^16^ databases via BLAST^36^. The extracted sequences were confirmed as orthologous to target NALCN/Cch1 subunits via reciprocal BLAST of the NCBI non-redundant database, and SmartBLAST^37^. Details about the general composition of these different sequence collections are provided in the results, and all sequences are available in FASTA format in Supplementary Files 2 (metazoan and fungal NALCN/Cch1), 3 (metazoan and fungal FAM155/Mid1), 4 (metazoan and fungal UNC80), 5 (fungal UNC80 only), 6 (metazoan and fungal UNC79), and 7 (fungal UNC79 only). Subsequently, these sequence sets were used as inputs for training subunit-specific custom HMM models using HMMER version 3.3.2, and the respective sequences identified with each model within the eukaryotic and FungiDB data sets were combined into corresponding sets and processes with CD-HIT^38^ using a sequence identity threshold of 99.9% to remove redundant sequences. The NALCN/Cch1 sequences were further processed by predicting transmembrane helices with Phobius^39^and discarding those with less than 18 transmembrane helices, to exclude single and tandem domain pore-loop channels from downstream analyses, and a preliminary cluster analysis with CLANS to remove sequences with less than three sequence similarity connections with other sequences. All final sets of identified NALCN/Cch1, FAM155, UNC79, and UNC80 sequences are provided in Supplementary Files 8 (NALCN/Cch1 homologues from eukaryotes), 9 (Cch1 homologues from Fungi), 10 (FAM155/Mid1 homologues from eukaryotes), 11 (Mid1 homologues from fungi), 12 (UNC80 homologues from eukaryotes), 13 (UNC80 homologues from fungi, 14 (UNC79 homologues from eukaryotes), and 15 (UNC79 homologues from fungi).

For NALCN subunit searches strictly within animals, most protein sequences were identified through BLAST searches of the NBCI non-redundant database or the NBCI transcriptome shotgun assembly (TSA) database using NALCN, FAM155, UNC79, and UNC80 protein sequences from humans, *C. elegans*, and *T. adhaerens* as queries. The exceptions are the identified *T. adhaerens* sequences which were extracted from a whole animal mRNA transcriptome^40^, the *Ptychodera flava* (hemichordate) NALCN and FAM155 sequences which were extracted from a gene model database derived from a genome sequencing effort^41^, and the UNC79 sequence from the sponge *A. queenslandica* which was fragmented on NCBI and was hence extracted from an available transcriptome assembly^42^. Candidate sequences identified with BLAST were verified as orthologous through reciprocal BLAST of the NCBI non-redundant database as well as SmartBLAST. All verified sequences used for downstream analysis are provided in Supplementary Files 16 (NALCN), 17 (FAM155), 18 (UNC80), and 19 (UNC79).

### Cluster analysis and phylogenetic inference

The various output files from our HMM searches through the eukaryotic proteome set (Supplementary Files 8, 10, 12, and 14) were analyzed with the BLAST-based all-against-all clustering algorithm CLuster ANalysis of Sequences (CLANS)^43^, using the following amino acid substitution matrices and expect-value cutoffs: NALCN/Cch1 - PAM30, 1E-20; FAM155/Mid1 - BLOSUM45, 1E-6; UNC79 - BLOSUM65, 1E-5; UNC80 - BLOSUM65, 1E-5. These various substitution matrices and expect value cutoffs were selected by comparing CLANS outputs from different parameters and selecting those that produced the best clustering. All cluster diagrams were annotated with the CLANS graphical user interface and exported as SVG files, and these were further annotated with Adobe Illustrator 2025 for generating figures.

For phylogenetic analyses, the various output files from our HMM and reciprocal BLAST searches (*i.e.*, Supplementary Files 8 to 19) were first aligned with the program MAFFT version 7.490^44^, then trimmed with trimAl^45^ using gappyout mode. These trimmed alignments were then used as input for the maximum likelihood inference algorithm IQ-TREE2^46^, using the ModelFinder^47^ option to identify the best-fit model for phylogenetic inference under the Bayesian Information Criterion. This resulted in the following best-fit models: NALCN/Cch1 in eukaryotes - Q.pfam+R10; Cch1 in fungi - Q.yeast+F+I+R8; NALCN in metazoans - Q.yeast+F+I+G4; FAM155/Mid1 in eukaryotes - VT+I+G4; Mid1 in fungi - Q.pfam+R8; FAM155 in metazoans - VT+R3; UNC80 in eukaryotes and fungi - Q.insect+F+I+R6; UNC80 in fungi - Q.yeast+F+I+G4; UNC80 in metazoans - Q.insect+F+R4; UNC79 in eukaryotes and fungi - Q.insect+R6; UNC79 in fungi - Q.insect+F+I+G4; UNC79 in metazoans - Q.insect+R4. For all trees, node support was estimated via 1000 replicate Shimodaira-Hasegawa approximate likelihood ratio tests (SH-aLRT)^48^, approximate Bayes tests (aBayes)^49^, and fast local bootstrap probability tests (LBP)^50^. Resulting phylogenetic trees were annotated with the programs FigTree version 1.4.4 and the Interactive Tree Of Life (iTOL) version 7^51^. Further annotation of the trees and final figure preparations were done using Adobe Illustrator 2025. Raw phylogenetic trees, in nexus format, are provided in Supplementary Files 20 (Figure 2A), 21 (Figure 2B), 22 (Figure 3B), 23 (Figure 3C), 24 (Figure 5B), 25 (Figure 5C), 26 (Figure 6B), 27 (Figure 6C), 28 (Figure S1), 29 (Figure S2), 30 (Figure S4), and 31 (Figure S5).

### Structure prediction and analysis

All structural predictions were done using AlphaFold3^18^. Corresponding sequence accession numbers, along with predicted template modeling (pTM) and interface predicted template modeling (ipTM) scores are provided in Supplementary Table 2. Structures were analyzed and visualized with the program ChimeraX version 1.9^52^, and images for figures were exported as.png files and further annotated with Adobe Illustrator 2025. Multiple sequence alignments were done with Clustal Omega^53^ and annotated with Excel and Adobe Illustrator.

## Contributions

AS, AM and PL conceived the project. AS, L-YG, BB, and WE conducted the sequence identification, and AS conducted the phylogenetic and clustering analyses. AS and TM conducted the structural analyses and sequence alignments. AS wrote the first draft of the manuscript and all authors contributed to revisions.

---

**Figure S1.**
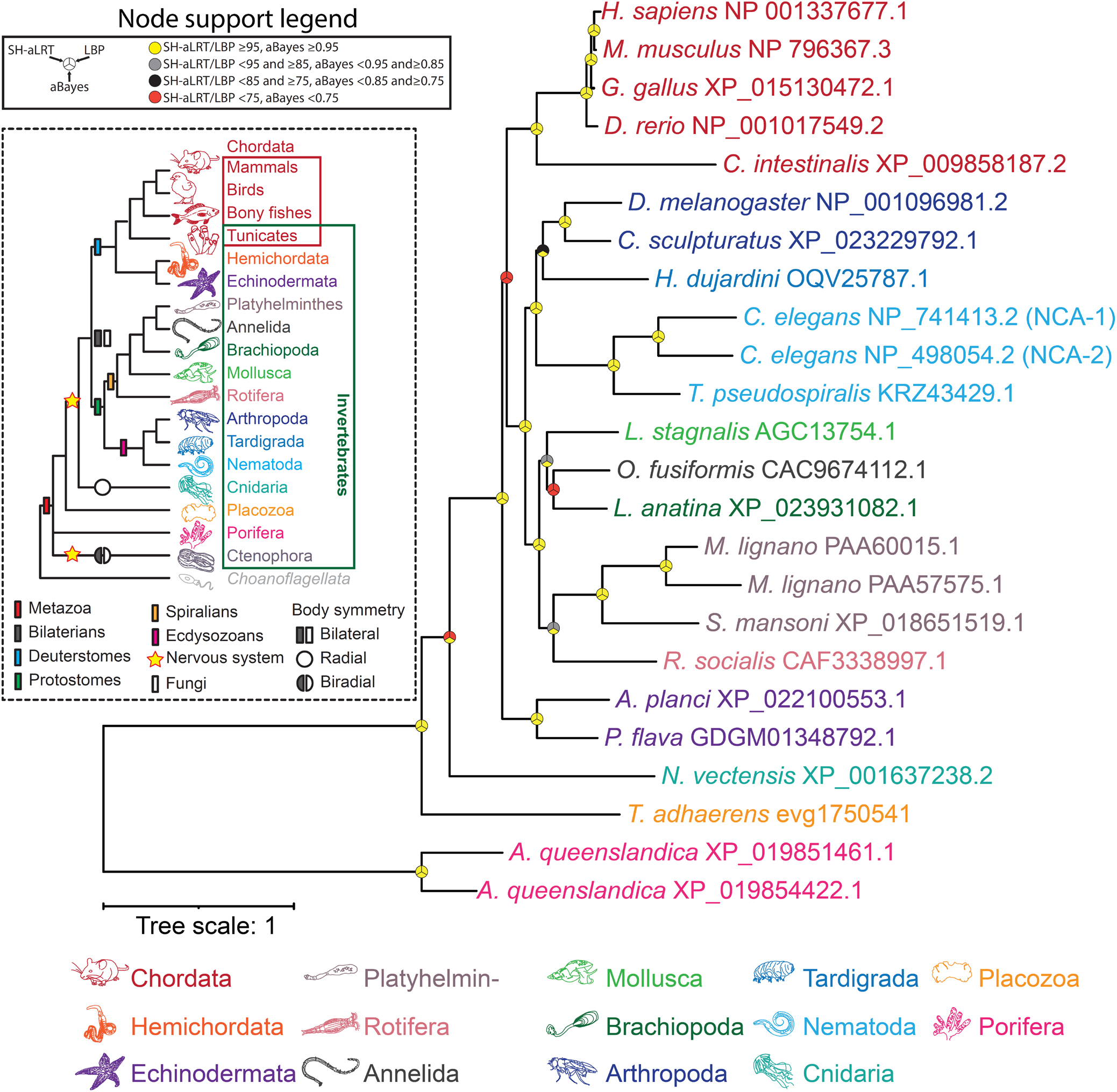
Maximum likelihood tree of manually identified NALCN homologues from animals. Node support values for three separate analyses, SH-aLRT, LBP, and aBayes, are depicted by circular symbols with colors reflecting ranges of values as indicated in the legend.

**Figure S2.**
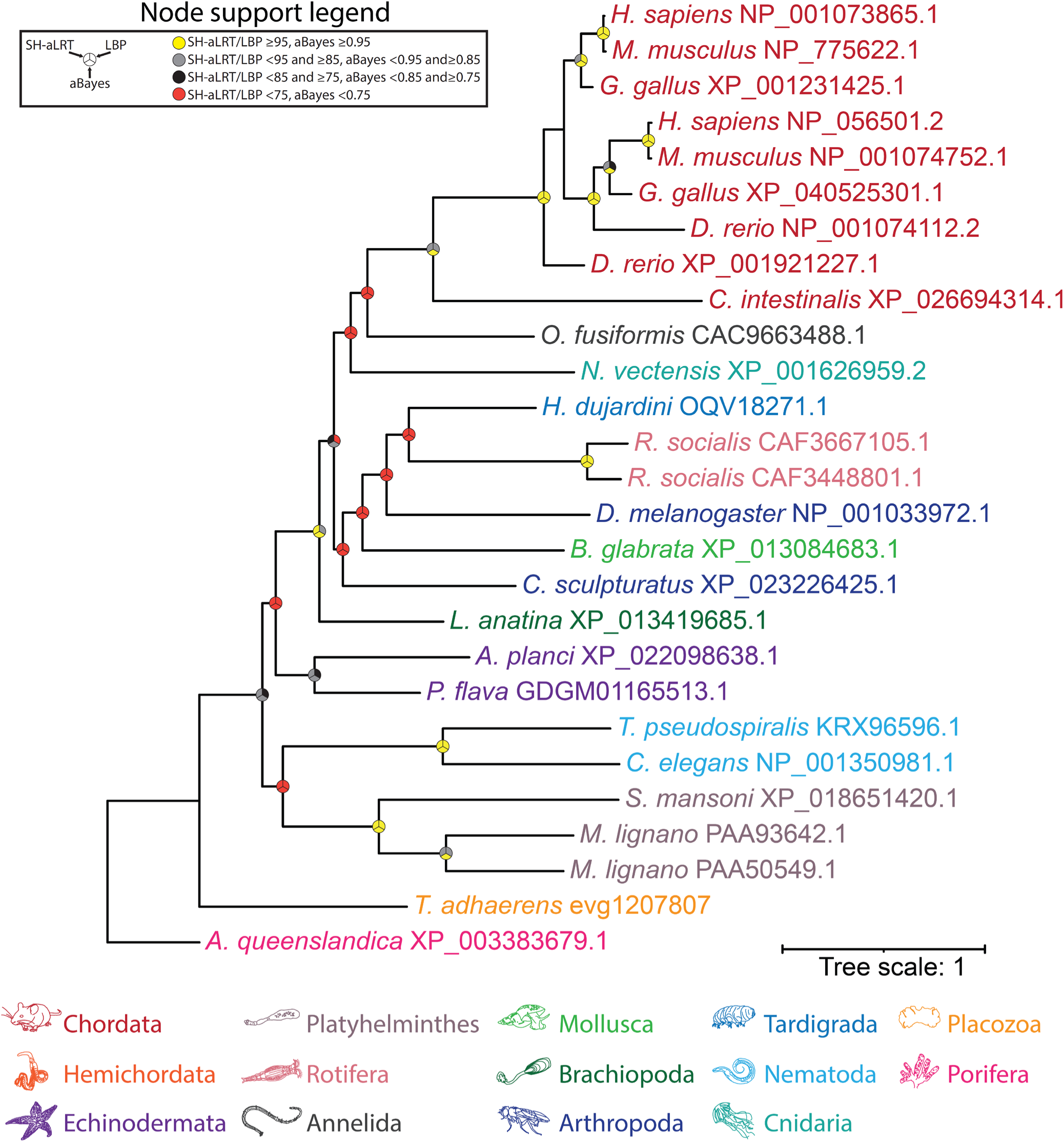
Maximum likelihood tree of manually identified FAM155 homologues from animals. Node support values for three separate analyses, SH-aLRT, LBP, and aBayes, are depicted by circular symbols with colors reflecting ranges of values as indicated in the legend.

**Figure S3.**
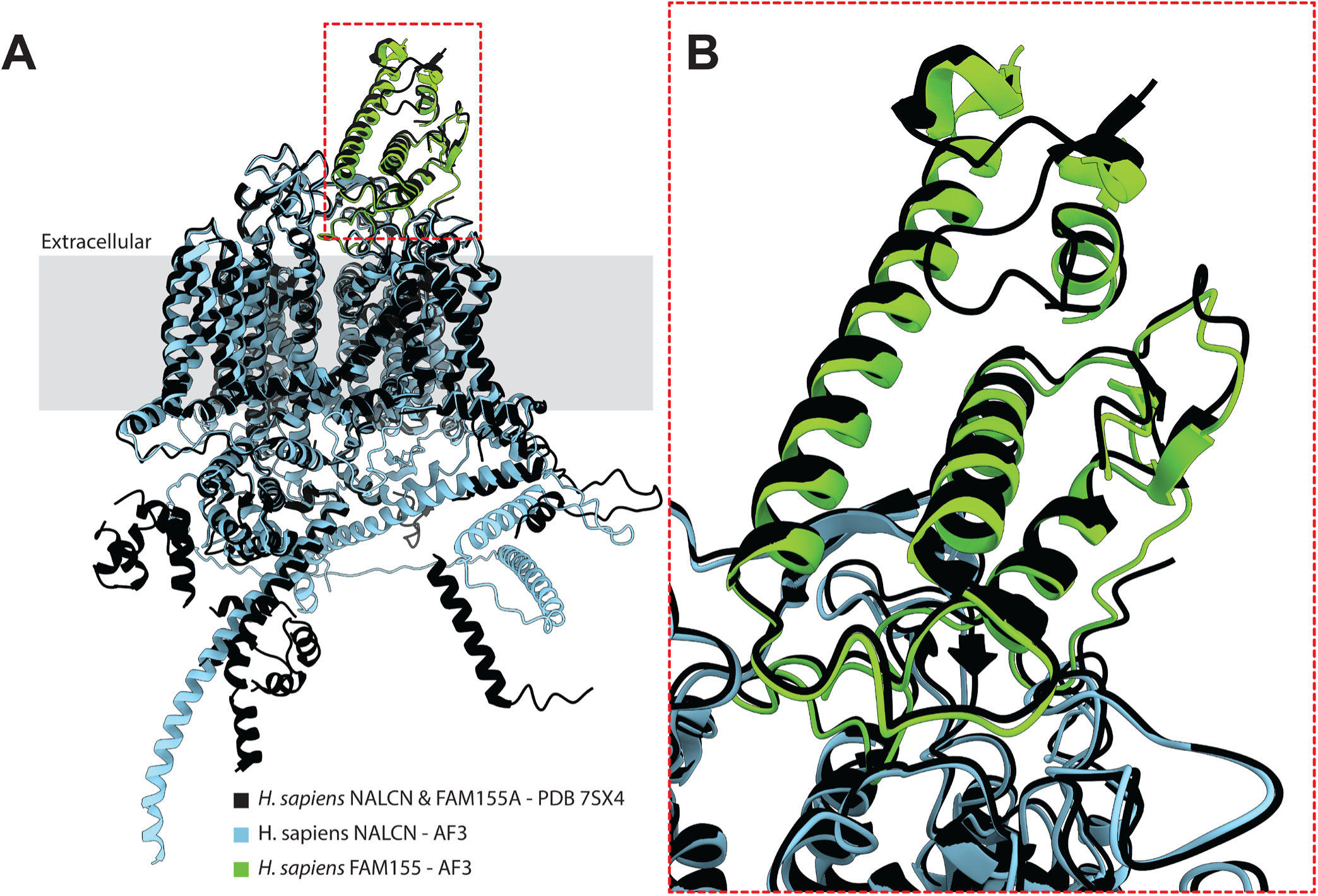
A) Structural alignment of the solved structure of human NALCN and FAM155A (from PDB accession number 7SX4) and AlphaFold3-predicted structures of these proteins in complex with each other. **B)** Close up view of the region corresponding to the red dashed box in panel A.

**Figure S4.**
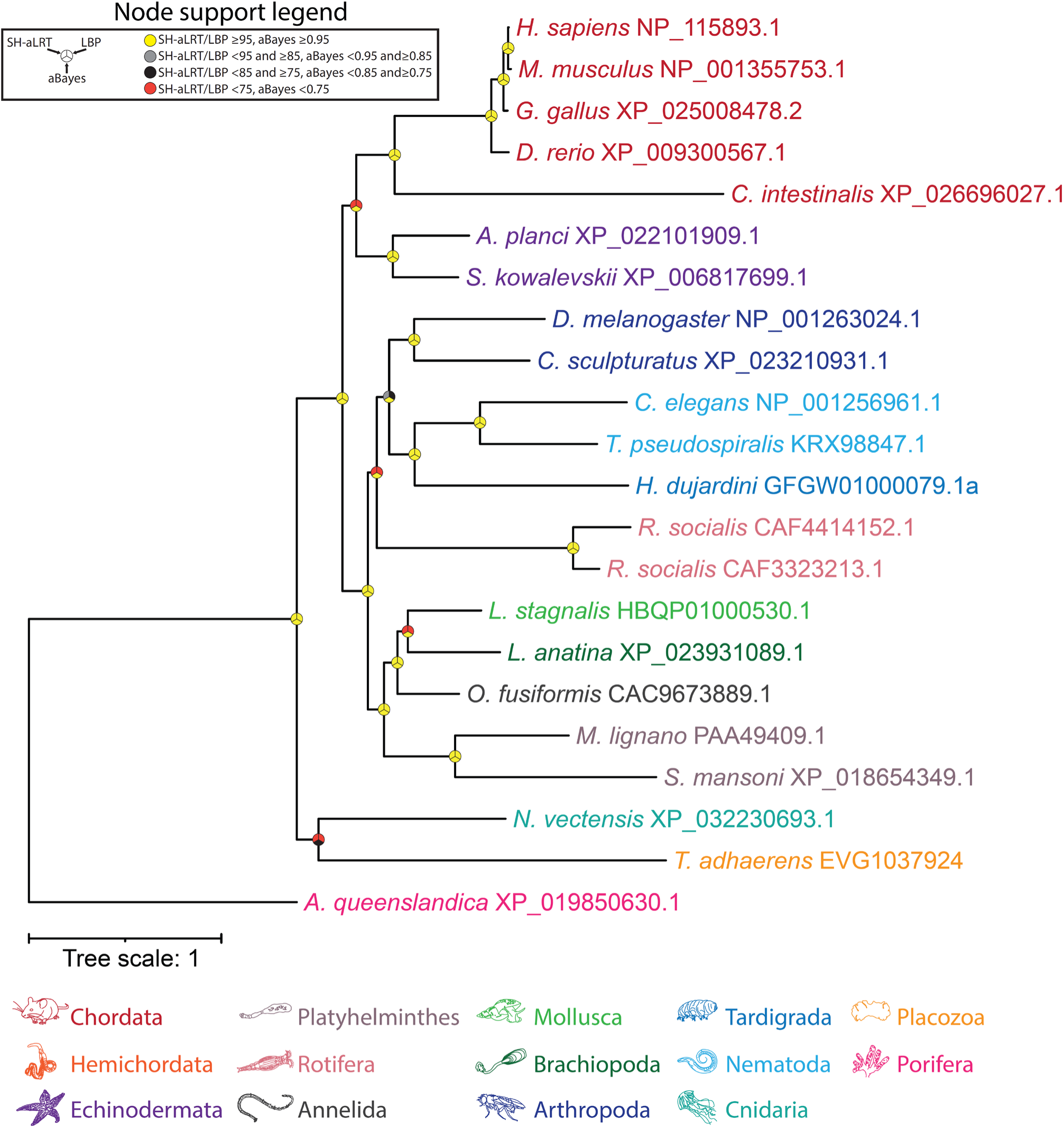
Maximum likelihood tree of manually identified UNC79 homologues from animals. Node support values for three separate analyses, SH-aLRT, LBP, and aBayes, are depicted by circular symbols with colors reflecting ranges of values as indicated in the legend.

**Figure S5.**
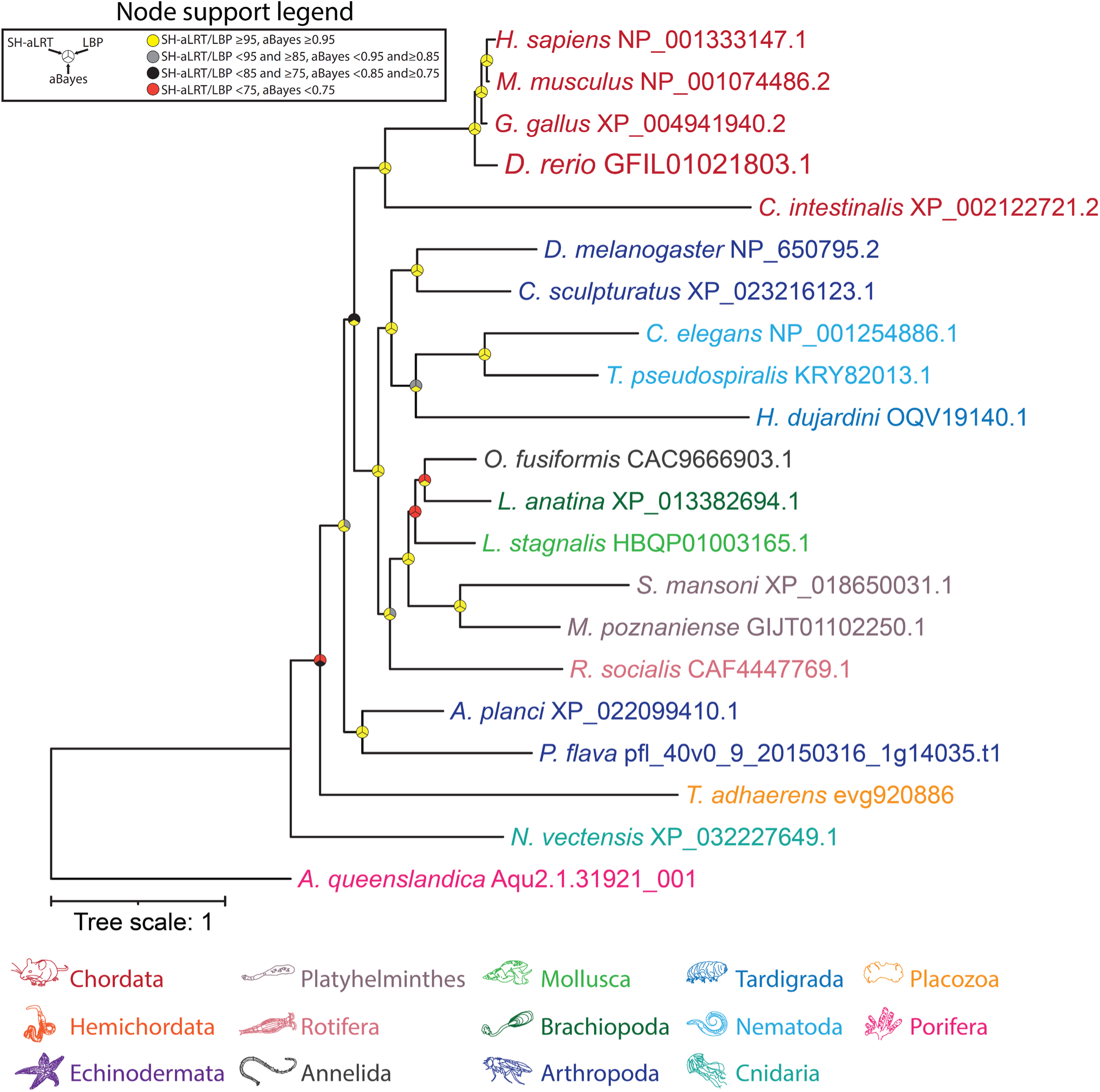
Maximum likelihood tree of manually identified UNC80 homologues from animals. Node support values for three separate analyses, SH-aLRT, LBP, and aBayes, are depicted by circular symbols with colors reflecting ranges of values as indicated in the legend.

## References

1 Liebeskind, B. J., Hillis, D. M. & Zakon, H. H. Phylogeny unites animal sodium leak channels with fungal calcium channels in an ancient, voltage-insensitive clade. Molecular biology and evolution 29, 3613–3616 (2012).

2 Pozdnyakov, I., Matantseva, O. & Skarlato, S. Diversity and evolution of four-domain voltage-gated cation channels of eukaryotes and their ancestral functional determinants. Scientific reports 8, 3539 (2018).

3 Monteil, A. et al. New insights into the physiology and pathophysiology of the atypical sodium leak channel NALCN. Physiological Reviews 104, 399–472 (2024).

4 Cochet-Bissuel, M., Lory, P. & Monteil, A. The sodium leak channel, NALCN, in health and disease. Frontiers in cellular neuroscience 8, 132 (2014).

5 Ren, D. Sodium leak channels in neuronal excitability and rhythmic behaviors. Neuron 72, 899–911 (2011).

6 Kang, Y. & Chen, L. Structure and mechanism of NALCN-FAM155A-UNC79-UNC80 channel complex. Nature Communications 13, 2639 (2022).

7 Kschonsak, M. et al. Structural architecture of the human NALCN channelosome. Nature 603, 180–186 (2022).

8 Zhou, L., Liu, H., Zhao, Q., Wu, J. & Yan, Z. Architecture of the human NALCN channelosome. Cell Discovery 8, 33 (2022).

9 Ghezzi, A., Liebeskind, B. J., Thompson, A., Atkinson, N. S. & Zakon, H. H. Ancient association between cation leak channels and Mid1 proteins is conserved in fungi and animals. Frontiers in molecular neuroscience 7, 15 (2014).

10 Peiter, E., Fischer, M., Sidaway, K., Roberts, S. K. & Sanders, D. The Saccharomyces cerevisiae Ca2+ channel Cch1pMid1p is essential for tolerance to cold stress and iron toxicity. FEBS letters 579, 5697–5703 (2005).

11 Vu, K., Bautos, J. M. & Gelli, A. The Cch1-Mid1 high-affinity calcium channel contributes to the virulence of Cryptococcus neoformans by mitigating oxidative stress. Eukaryotic cell 14, 1135–1143 (2015).

12 Yu, Q. et al. Roles of Cch1 and Mid1 in morphogenesis, oxidative stress response and virulence in Candida albicans. Mycopathologia 174, 359–369 (2012).

13 Harren, K. & Tudzynski, B. Cch1 and Mid1 are functionally required for vegetative growth under low-calcium conditions in the phytopathogenic ascomycete Botrytis cinerea. Eukaryotic Cell 12, 712–724 (2013).

14 De Castro, P. A. et al. The involvement of the Mid1/Cch1/Yvc1 calcium channels in Aspergillus fumigatus virulence. PLoS One 9, e103957 (2014).

15 Madera, M. & Gough, J. A comparison of profile hidden Markov model procedures for remote homology detection. Nucleic acids research 30, 4321–4328 (2002).

16 Basenko, E. Y. et al. FungiDB: an integrated bioinformatic resource for fungi and oomycetes. Journal of Fungi 4, 39 (2018).

17 Li, Y. et al. A genome-scale phylogeny of the kingdom Fungi. Current Biology 31, 1653–1665. e1655 (2021).

18 Abramson, J. et al. Accurate structure prediction of biomolecular interactions with AlphaFold 3. Nature 630, 493–500 (2024).

19 Liu, M., Du, P., Heinrich, G., Cox, G. M. & Gelli, A. Cch1 mediates calcium entry in Cryptococcus neoformans and is essential in low-calcium environments. Eukaryotic Cell 5, 1788–1796 (2006).

20 Hong, M.-P. et al. Activity of the calcium channel pore Cch1 is dependent on a modulatory region of the subunit Mid1 in Cryptococcus neoformans. Eukaryotic cell 12, 142–150 (2013).

21 Brown, M. W. et al. Phylogenomics places orphan protistan lineages in a novel eukaryotic super-group. Genome Biology and Evolution 10, 427–433 (2018).

22 Kschonsak, M. et al. Structure of the human sodium leak channel NALCN. Nature 587, 313–318 (2020).

23 Xie, J. et al. Structure of the human sodium leak channel NALCN in complex with FAM155A. Nature communications 11, 5831 (2020).

24 Kang, Y., Wu, J.-X. & Chen, L. Structure of voltage-modulated sodium-selective NALCN- FAM155A channel complex. Nature communications 11, 6199 (2020).

25. 25 Boucher, M. J., et al. Phenotypic landscape of a fungal meningitis pathogen reveals its unique biology. *bioRxiv* (2024).

26 Tewari, R., Bailes, E., Bunting, K. A. & Coates, J. C. Armadillo-repeat protein functions: questions for little creatures. Trends in cell biology 20, 470–481 (2010).

27 Schultz, D. T. et al. Ancient gene linkages support ctenophores as sister to other animals. Nature 618, 110–117 (2023).

28 Dunn, C. W. et al. Broad phylogenomic sampling improves resolution of the animal tree of life. Nature 452, 745–749 (2008).

29 McCleskey, E. W. Calcium channel permeation: a field in flux. Journal of General Physiology 113, 765–772 (1999).

30 Talavera, K. & Nilius, B. Biophysics and structure–function relationship of T-type Ca2+ channels. Cell calcium 40, 97–114 (2006).

31 Barzilai, M. G. et al. Convergent evolution of sodium ion selectivity in metazoan neuronal signaling. Cell reports 2, 242–248 (2012).

32 Yang, J., Elllnor, P. T., Sather, W. A., Zhang, J.-F. & Tsien, R. W. Molecular determinants of Ca2+ selectivity and ion permeation in L-type Ca2+ channels. Nature 366, 158–161 (1993).

33 Senatore, A., Monteil, A., van Minnen, J., Smit, A. B. & Spafford, J. D. NALCN ion channels have alternative selectivity filters resembling calcium channels or sodium channels. PLoS One 8, e55088 (2013).

34 Simão, F. A., Waterhouse, R. M., Ioannidis, P., Kriventseva, E. V. & Zdobnov, E. M. BUSCO: assessing genome assembly and annotation completeness with single-copy orthologs. Bioinformatics 31, 3210–3212 (2015).

35 Sayers, E. W. et al. Database resources of the national center for biotechnology information. Nucleic acids research 49, D10–D17 (2021).

36 Altschul, S. F., Gish, W., Miller, W., Myers, E. W. & Lipman, D. J. Basic local alignment search tool. Journal of molecular biology 215, 403–410 (1990).

37 Database resources of the national center for biotechnology information. Nucleic acids research 46, D8-D13 (2018).

38 Li, W. & Godzik, A. Cd-hit: a fast program for clustering and comparing large sets of protein or nucleotide sequences. Bioinformatics 22, 1658–1659 (2006).

39 Käll, L., Krogh, A. & Sonnhammer, E. L. A combined transmembrane topology and signal peptide prediction method. Journal of molecular biology 338, 1027–1036 (2004).

40 Wong, Y. Y., Le, P., Elkhatib, W., Piekut, T. & Senatore, A. Transcriptome profiling of Trichoplax adhaerens highlights its digestive epithelium and a rich set of genes for fast electrogenic and slow neuromodulatory cellular signaling. (2019).

41 Simakov, O. et al. Hemichordate genomes and deuterostome origins. Nature 527, 459–465 (2015).

42 Fernandez-Valverde, S. L., Calcino, A. D. & Degnan, B. M. Deep developmental transcriptome sequencing uncovers numerous new genes and enhances gene annotation in the sponge Amphimedon queenslandica. BMC genomics 16, 1–11 (2015).

43 Zimmermann, L. et al. A completely reimplemented MPI bioinformatics toolkit with a new HHpred server at its core. Journal of molecular biology 430, 2237–2243 (2018).

44 Katoh, K. & Standley, D. M. MAFFT multiple sequence alignment software version 7: improvements in performance and usability. Molecular biology and evolution 30, 772–780 (2013).

45 Capella-Gutiérrez, S., Silla-Martínez, J. M. & Gabaldón, T. trimAl: a tool for automated alignment trimming in large-scale phylogenetic analyses. Bioinformatics 25, 1972–1973 (2009).

46 Minh, B. Q. et al. IQ-TREE 2: new models and efficient methods for phylogenetic inference in the genomic era. Molecular biology and evolution 37, 1530–1534 (2020).

47 Kalyaanamoorthy, S., Minh, B. Q., Wong, T. K., Von Haeseler, A. & Jermiin, L. S. ModelFinder: fast model selection for accurate phylogenetic estimates. Nature methods 14, 587–589 (2017).

48 Guindon, S. et al. New algorithms and methods to estimate maximum-likelihood phylogenies: assessing the performance of PhyML 3.0. Systematic biology 59, 307–321 (2010).

49 Anisimova, M., Gil, M., Dufayard, J.-F., Dessimoz, C. & Gascuel, O. Survey of branch support methods demonstrates accuracy, power, and robustness of fast likelihood-based approximation schemes. Systematic biology 60, 685–699 (2011).

50 Adachi, J. & Hasegawa, M. MOLPHY version 2.3: programs for molecular phylogenetics based on maximum likelihood. Vol. 28 (Institute of Statistical Mathematics Tokyo, 1996).

51 Letunic, I. & Bork, P. Interactive Tree Of Life (iTOL): an online tool for phylogenetic tree display and annotation. Bioinformatics 23, 127–128 (2007).

52 Meng, E. C. et al. UCSF ChimeraX: Tools for structure building and analysis. Protein Science 32, e4792 (2023).

53 Sievers, F. et al. Fast, scalable generation of high-quality protein multiple sequence alignments using Clustal Omega. Molecular systems biology 7, 539 (2011).

54 Al Jewari, C. & Baldauf, S. L. An excavate root for the eukaryote tree of life. Science Advances 9, eade4973 (2023).

55 Strassert, J. F., Irisarri, I., Williams, T. A. & Burki, F. A molecular timescale for eukaryote evolution with implications for the origin of red algal-derived plastids. Nature Communications 12, 1879 (2021).

