## Supplementary Files 1 to 31 for "NALCN/Cch1 channelosome subunits originated in early eukaryotes and are fully conserved in animals, fungi, and apusomonads": File S1 - Aligments of human and fungal UNC79 and UNC80.docx

```
UNC80
>KAG1050648.1 hypothetical protein G6F43_007098 [Rhizopus delemar]
MSQEDQNDLVKNYKEESSRDATLDKQYKKKTKHFSAWDKLRENVSEYSSDTLYHATSDPSRPNMSKMSSPMHKKQMYNPFLRNQHTTTAENPLHDKINLTNTTLVAAQAAGMSSFVRNFKSKVGAGRHNGAIYATSTAVQQDIYRLERDLEKLLSHMNNRSQNASFTESESSNHNTVYSSDSLALNNFSVKSASKKTIFHRHSQAVLDPSTQHNEKHLDVTRISANITSIMDIVNKHKSATRLPFTSEILAVLSIPFDRKPQKTEDCIQALDAFDHIRSRFTLLESSEHFEQVLFCCSILGSSGIDLKEKILDILKKLVEPCFTYPNYLPSTPTAFHALAYALATSFAKQTQNIELQEKKTTYDLVRSGILSLLDNLANGKLISVQGDQWTTYFHTTQAASASLAKICMVESLCKCLMVSICRQTSFHERNAELFLNNLIQKDQDIPTLLELLQRYWTEPDTEILPAYFKSLGLLTELASEISLALSIEKLKSSAIPLLFNFILEKSAPSNVYNYLNQAISGKIQKSVIFNIITMLLCFLSVDLLTAPSRSAPSSNTPRTPPNLTPQSSLNDMFISSPVLDDSGSEAPHIIVSPEEAFLSQLLETAKAYLIEMWNYGYKSHICKCIEIMLQDTSENQLAHTYQNLLFHIDIAIGEEIAKVTIPILFKRILDTSPAPTPDFCSMLYQLSQRFRPLFYKPVVSCVASNDNDKVASLLKLMTSLRQYLSGVQFWMQDAEMINVLLLSDVGPKKLQRETSSLQQNHTSSTNLLTWGSTTLGQCIIATELVWVLKELREKQKDPDRNMEEDEVAKKFLIDLERRLALFLTAKEKKVLVPLPLRIILCNLFADARFFCYTTHRPGWLTCAINWATQPRTNDIGSQAISETFLDENLSETGFCSNDMSLPVLNANHIEEALIMFQRMQNTYLDVVEYFRDESSTSHGDSHGTRQAASSLLFSNTQMTENLKYGRRHLAETMYPINQSSSAALNLNPPEFDHLENNEGESNSIMLDLAKHKFDGIEEINQDPFGSVFSLLAAVYSTLSSHELGRLLKPLWKTYMDDKKAASFTPAAFLLMECGEKIPKLTTEMFSQALQRLNILNQSYIQISSRNKPFRGDGGAFATPFLPTDLGSNEFSLDEPRWMSKLKNTSNLPIELKRQIQELGWNEDDNAEEQEAFKKALTPLGLLPSLFLKDDNEDTNDADNPNIVVIRERGKIVDISKIITRRKRTATVHSFTAMFLTTVDLLMDDFAGVNNVLRELIEQFLKDDPALFLRAFLNNLGKHKADSLKDTLTRIRYIISMQRKLPPGLTHILFNYLAGMLKWIVRENQPDGLKFMTLVHPILAELTLSTNDLSVRDLRKNKIEHLLVSTGKFWFTHEQPADMFPRYLSNEQHSFAVLNIPQQIFFVACLRISHIQFLINYLARFPREVYAVKKTLQDYEPMPVPGSKWETRQLMEKAYYPDLSRRKNRQSNHKQEPLDTEDSQQHFDKEMLSLMRARIWLRFIDVLLNGLNKNYNDHDELERILQGVNTIMIEHSHDFGIIGQILILYTRMVTKFKRLFKSNRGYIIFLPALFKVFCESEDNPQITSAITFAWCRFYAVHEEAFVFQMLGALVPNILMIYGQSDEVGHWMIENLFKLMQAMDNPPHLGSAPDTLGLQLQVELDDHERSIQDRIDTASSQVSSALSVSIFKPLARGVTAPISQMEISTYNNRPFKLEDFIKLFLTVIAYDPGSLRAEQFVKVFTFLIPFFLEQSHLKSMMHEGASALVNVFVKFSKSVKSVVTTNSHTTSGEYQGISSSDSRLPGSNNAANTGPKAESAQNAYGKQWQQNDRFTIKQEFITLVHKYMKHGGTLSEVSHEKMAQIIRIVLRDYRNIKGLTVKTDWIKDYLVVSMYSMVDTRNYTKSLKKVLSEIYAQYRVQWKTVDASDLYEGLAIMLEQGQGKAIMMNDLAGLVKEKFVSFGLTMATLTDWQNNTKGQVKFCNSLVRLIIAILENSSQDVFAEIERQVPSVGLIGRVVIPLCLQYNLQWEYSTVGISSTSELNSANNWTRLLNFIAGVCSHTSLLKSKSSGFSLSQFTTAMGQTNIEDEVDMSDFPEAKDGKQTPQSVAVLFSLSLIAIKIILIRSHNTFNDLKGSWSQIAFFIRNTLVFGQTLKLLKAKVGRSDSYNLASPSDVLSPGMPSPSFFSPLGLSSPGASPYLSSNNPSLSPDLRANFSSNAPLGTGTIYDFTTWRFLEFVVCFRTPLFVLLKDFIHEKLGQMGPNVGSYRNSLYSPRASFNFPQNENTRWKSWGAEHSSVNTSGDGQSNANASAHSIVIPKLNIEDTGGVDAETLPTFGLGLHKSNSSASPGHVSHSPTGYSETDSSSPRSPTTNPFYKQQQHMSPGADVSKENNLHLLQAETITSVVNVQVAVGFKPALPWITMDSRDRSKPWNKKEAVARIANEWQLLLQLSTETDPLVNKANNNGSSANISASPLV


>Homo_sapiens_NP_115893.1
MVKRKSSEGQEQDGGRGIPLPIQTFLWRQTSAFLRPKLGKQYEASCVSFERVLVENKLHGLSPALSEAIQSISRWELVQAALPHVLHCTATLLSNRNKLGHQDKLGVAETKLLHTLHWMLLEAPQDCNNERFGGTDRGSSWGGSSSAFIHQVENQGSPGQPCQSSSNDEEENNRRKIFQNSMATVELFVFLFAPLVHRIKESDLTFRLASGLVIWQPMWEHRQPGVSGFTALVKPIRNIITAKRSSPINSQSRTCESPNQDARHLEGLQVVCETFQSDSISPKATISGCHRGNSFDGSLSSQTSQERGPSHSRASLVIPPCQRSRYATYFDVAVLRCLLQPHWSEEGTQWSLMYYLQRLRHMLEEKPEKPPEPDIPLLPRPRSSSMVAAAPSLVNTHKTQDLTMKCNEEEKSLSSEAFSKVSLTNLRRSAVPDLSSDLGMNIFKKFKSRKEDRERKGSIPFHHTGKRRPRRMGVPFLLHEDHLDVSPTRSTFSFGSFSGLGEDRRGIEKGGWQTTILGKLTRRGSSDAATEMESLSARHSHSHHTLVSDLPDPSNSHGENTVKEVRSQISTITVATFNTTLASFNVGYADFFNEHMRKLCNQVPIPEMPHEPLACANLPRSLTDSCINYSYLEDTEHIDGTNNFVHKNGMLDLSVVLKAVYLVLNHDISSRICDVALNIVECLLQLGVVPCVEKNRKKSENKENETLEKRPSEGAFQFKGVSGSSTCGFGGPAVSGAGDGGGEEGGGGDGGGGGGDGGGGGGGGGGPYEKNDKNQEKDESTPVSNHRLALTMLIKIVKSLGCAYGCGEGHRGLSGDRLRHQVFRENAQNCLTKLYKLDKMQFRQTMRDYVNKDSLNNVVDFLHALLGFCMEPVTDNKAGFGNNFTTVDNKSTAQNVEGIIVSAMFKSLITRCASTTHELHSPENLGLYCDIRQLVQFIKEAHGNVFRRVALSALLDSAEKLAPGKKVEENEQESKPAGSKRSEAGSIVDKGQVSSAPEECRSFMSGRPSQTPEHDEQMQGANLGRKDFWRKMFKSQSAASDTSSQSEQDTSECTTAHSGTTSDRRARSRSRRISLRKKLKLPIGKRNWLKRSSLSGLADGVEDLLDISSVDRLSFIRQSSKVKFTSAVKLSEGGPGSGMENGRDEEENFFKRLGCHSFDDHLSPNQDGGKSKNVVNLGAIRQGMKRFQFLLNCCEPGTIPDASILAAALDLEAPVVARAALFLECARFVHRCNRGNWPEWMKGHHVNITKKGLSRGRSPIVGNKRNQKLQWNAAKLFYQWGDAIGVRLNELCHGESESPANLLGLIYDEETKRRLRKEDEEEDFLDDSTVNPSKCGCPFALKMAACQLLLEITTFLRETFSCLPRPRTEPLVDLESCRLRLDPELDRHRYERKISFAGVLDENEDSKDSLHSSSHTLKSDAGVEEKKEGSPWSASEPSIEPEGMSNAGAEENYHRNMSWLHVMILLCNQQSFICTHVDYCHPHCYLHHSRSCARLVRAIKLLYGDSVDSLRESSNISSVALRGKKQKECSDKSCLRTPSLKKRVSDANLEGKKDSGMLKYIRLQVMSLSPAPLSLLIKAAPILTEEMYGDIQPAAWELLLSMDEHMAGAAAAMFLLCAVKVPEAVSDMLMSEFHHPETVQRLNAVLKFHTLWRFRYQVWPRMEEGAQQIFKIPPPSINFTLPSPVLGMPSVPMFDPPWVPQCSGSVQDPINEDQSKSFSARAVSRSHQRAEHILKNLQQEEEKKRLGREASLITAIPITQEACYEPTCTPNSEPEEEVEEVTNLASRRLSVSPSCTSSTSHRNYSFRRGSVWSVRSAVSAEDEEHTTEHTPNHHVPQPPQAVFPACICAAVLPIVHLMEDGEVREDGVAVSAVAQQVLWNCLIEDPSTVLRHFLEKLTISNRQDELMYMLRKLLLNIGDFPAQTSHILFNYLVGLIMYFVRTPCEWGMDAISATLTFLWEVVGYVEGLFFKDLKQTMKKEQCEVKLLVTASMPGTKTLVVHGQNECDIPTQLPVHEDTQFEALLKECLEFFNIPESQSTHYFLMDKRWNLIHYNKTYVRDIYPFRRSVSPQLNLVHMHPEKGQELIQKQVFTRKLEEVGRVLFLISLTQKIPTAHKQSHVSMLQEDLLRLPSFPRSAIDAEFSLFSDPQAGKELFGLDTLQKSLWIQLLEEMFLGMPSEFPWGDEIMLFLNVFNGALILHPEDSALLRQYAATVINTAVHFNHLFSLSGYQWILPTMLQVYSDYESNPQLRQAIEFACHQFYILHRKPFVLQLFASVAPLLEFPDAANNGPSKGVSAQCLFDLLQSLEGETTDILDILELVKAEKPLKSLDFCYGNEDLTFSISEAIKLCVTVVAYAPESFRSLQMLMVLEALVPCYLQKLKRQTSQVETVPAAREEIAATAALATSLQALLYSVEVLTRPMTAPQMSRCDQGHKGTTTANHTMSSGVNTRYQEQGAKLHFIRENLHLLEEGQGIPREELDERIAREEFRRPRESLLNICTEFYKHCGPRLKILQNLAGEPRVIALELLDVKSHMRLAEIAHSLLKLAPYDTQTMESRGLRRYIMEMLPITDWTAEAVRPALILILKRLDRMFNKIHKMPTLRRQVEWEPASNLIEGVCLTLQRQPIISFLPHLRSLINVCVNLVMGVVGPSSVADGLPLLHLSPYLSPPLPFSTAVVRLVALQIQALKEDFPLSHVISPFTNQERREGMLLNLLIPFVLTVGSGSKDSPWLEQPEVQLLLQTVINVLLPPRIISTSRSKNFMLESSPAHCSTPGDAGKDLRREGLAESTSQAAYLALKVILVCFERQLGSQWYWLSLQVKEMALRKVGGLALWDFLDFIVRTRIPIFVLLRPFIQCKLLAQPAENHEELSARQHIADQLERRFIPRPLCKSSLIAEFNSELKILKEAVHSGSAYQGKTSISTVGTSTSAYRLSLATMSRSNTGTGTVWEQDSEPSQQASQDTLSRTDEEDEENDSISMPSVVSEQEAYLLSAIGRRRFSSHVSSMSVPQAEVGMLPSQSEPNVLDDSQGLAAEGSLSRVASIQSEPGQQNLLVQQPLGRKRGLRQLRRPLLSRQKTQTEPRNRQGARLSTTRRSIQPKTKPSADQKRSVTFIEAQPEPAAAPTDALPATGQLQGCSPAPSRKPEAMDEPVLTSSPAIVVADLHSVSPKQSENFPTEEGEKEEDTEAQGATAHSPLSAQLSDPDDFTGLETSSLLQHGDTVLHISEENGMENPLLSSQFTFTPTELGKTDAVLDESHV


CLUSTAL format alignment by MAFFT (v7.511)


KAG1050648 M----SQEDQNDLVKNYKEESSRD------------------ATLDKQYKKKTKHF----
Homo_sapie MVKRKSSEGQ-------EQDGGRGIPLPIQTFLWRQTSAFLRPKLGKQYEASCVSFERVL
           *    *.*.*       :::..*.                  ..*.***: .   *

KAG1050648 -------------------SAW---------------------------DKLRENVSEY-
Homo_sapie VENKLHGLSPALSEAIQSISRWELVQAALPHVLHCTATLLSNRNKLGHQDKL--GVAETK
                              * *                           ***  .*:*

KAG1050648 --------------------------------SSDTLYHATSDPSRPNMSKMSS------
Homo_sapie LLHTLHWMLLEAPQDCNNERFGGTDRGSSWGGSSSAFIHQVENQGSPGQPCQSSSNDEEE
                                           **.:: * ..: . *. .  **

KAG1050648 ---------------------------------------------PM--HKKQMYNPF--
Homo_sapie NNRRKIFQNSMATVELFVFLFAPLVHRIKESDLTFRLASGLVIWQPMWEHRQPGVSGFTA
                                                        **  *::   . *

KAG1050648 ----LR-----------NQHTTTAENPLHDKINLTNTTLVAA--QAAGMS----------
Homo_sapie LVKPIRNIITAKRSSPINSQSRTCESPNQDARHLEGLQVVCETFQSDSISPKATISGCHR
               :*           *.:: *.*.* :*  :* .  :*.   *: .:*

KAG1050648 --SFVRNFKSKVGAGR---HNGAI----------YAT--STAVQQDI-------------
Homo_sapie GNSFDGSLSSQTSQERGPSHSRASLVIPPCQRSRYATYFDVAVLRCLLQPHWSEEGTQWS
             **  .:.*:..  *   *. *           ***  ..** : :

KAG1050648 --YRLERDLEKLLSHM---------------------------------NNRSQNASF--
Homo_sapie LMYYLQR-----LRHMLEEKPEKPPEPDIPLLPRPRSSSMVAAAPSLVNTHKTQDLTMKC
             * *:*     * **                                 .:::*: ::

KAG1050648 TESESSNHNTVYS---------------SDSLALN-------------------------
Homo_sapie NEEEKSLSSEAFSKVSLTNLRRSAVPDLSSDLGMNIFKKFKSRKEDRERKGSIPFHHTGK
           .*.*.*  . .:*               *..*.:*

KAG1050648 ------------------------NFSVKSAS--------------KKTIF---------
Homo_sapie RRPRRMGVPFLLHEDHLDVSPTRSTFSFGSFSGLGEDRRGIEKGGWQTTILGKLTRRGSS
                                   .**. * *              :.**:

KAG1050648 ------------HRHSQAVL-----DPSTQHNEKHLDVTRISANITSIM-----------
Homo_sapie DAATEMESLSARHSHSHHTLVSDLPDPSNSHGEN--TVKEVRSQISTITVATFNTTLASF
                       * **: .*     ***..*.*:   *..: ::*::*

KAG1050648 -----DIVNKH--KSATRLPFTS---EILAVLSIPFDRKPQKTEDCIQALDAFDHIRSRF
Homo_sapie NVGYADFFNEHMRKLCNQVPIPEMPHEPLACANLP----RSLTDSCIN-----------Y
                *:.*:*  * ..::*:..   * **  .:*     . *:.**:           :

KAG1050648 TLLESSEHFEQVLFCCSILGSSGI----------------DLKEKILDILKKLVE-----
Homo_sapie SYLEDTEHIDGT---NNFVHKNGMLDLSVVLKAVYLVLNHDISSRICDVALNIVECLLQL
           : **.:**:: .    .:: ..*:                *:..:* *:  ::**

KAG1050648 ---PC------------------------FTYPNYLPSTPTAFHALAYALATS-------
Homo_sapie GVVPCVEKNRKKSENKENETLEKRPSEGAFQFKGVSGSSTCGFGGPAVSGAGDGGGEEGG
              **                        * : .   *:. .* . * : * .

KAG1050648 ---------------------FAKQTQNIELQEKK--TTYDLVRSGILSLLDNL------
Homo_sapie GGDGGGGGGDGGGGGGGGGGPYEKNDKNQEKDESTPVSNHRLALTMLIKIVKSLGCAYGC
                                : *: :* * :*..  :.: *. : ::.::..*

KAG1050648 ANGKLISVQGDQW--TTYFHTTQAASASLAKICMVESLCKCLMVSICRQTSFHERNAELF
Homo_sapie GEGHR-GLSGDRLRHQVFRENAQNCLTKLYKLDKMQF----------RQT----------
           .:*:  .:.**:    .: ..:* . :.* *:  ::           ***

KAG1050648 LNNLIQKD--QDIPTLLELLQRYWTEPDTE------------------------ILPAYF
Homo_sapie MRDYVNKDSLNNVVDFLHALLGFCMEPVTDNKAGFGNNFTTVDNKSTAQNVEGIIVSAMF
           :.: ::**  :::  :*. *  :  ** *:                        *:.* *

KAG1050648 KSLGLLTELASEISLALSIEKLK-SSAIPLLFNFILEKSAPSNVYNYLNQAISGKIQKSV
Homo_sapie KS--LITRCASTTHELHSPENLGLYCDIRQLVQFI--KEAHGNVFRRV--ALSA------
           **  *:*. **      * *:*   . *  *.:**  *.* .**:. :  *:*.

KAG1050648 IFNIITMLLCFLSVDLLTAPSRSAPSSNTPRTPPNLTPQSSLNDMFISSPVLDDSGSEAP
Homo_sapie ---------------LLDSAEKLAPGKKVEENEQESKPAGS-------------KRSEAG
                          ** :..: **..:. ..  : .* .*             . ***

KAG1050648 HII------VSPEE--AFLS----------QLLETAKAYLIEMWNYGYKSH--ICKCIEI
Homo_sapie SIVDKGQVSSAPEECRSFMSGRPSQTPEHDEQMQGANLGRKDFWRKMFKSQSAASDTSSQ
            *:       :***  :*:*          : :: *:    ::*.  :**:   ..  .

KAG1050648 MLQDTSENQLAHTYQ-------------NLLFHIDIAIGEE--IAKVTIPILFKRILDTS
Homo_sapie SEQDTSECTTAHSGTTSDRRARSRSRRISLRKKLKLPIGKRNWLKRSSLSGLADGVEDLL
             *****   **:               .*  ::.:.**:.  : : ::. * . : *

KAG1050648 PAPTPDFCSMLYQLSQ-RFRPL---------------------FYKPVVSCVA-------
Homo_sapie DISSVDRLSFIRQSSKVKFTSAVKLSEGGPGSGMENGRDEEENFFKR-LGCHSFDDHLSP
             .: *  *:: * *: :* .                      *:*  :.* :

KAG1050648 SNDNDKVASLLKLMTSLRQYLSGVQFWMQ--------DAEMI-------------NVLLL
Homo_sapie NQDGGKSKNVVNL-GAIRQGMKRFQFLLNCCEPGTIPDASILAAALDLEAPVVARAALFL
           .:*..*  .:::*  ::** :. .** ::        **.::              .*:*

KAG1050648 S----------------------DVGPKKLQRETSSLQQNHTSSTNLLTWGSTTL-----
Homo_sapie ECARFVHRCNRGNWPEWMKGHHVNITKKGLSRGRSPIVGNKRNQK--LQWNAAKLFYQWG
           .                      ::  * *.*  *.:  *: ...  * *.::.*

KAG1050648 ------------GQCIIATELVWVLKELREKQKDPDRNMEEDEVAKKFLIDLERRLALFL
Homo_sapie DAIGVRLNELCHGESESPANLLGLIYDEETKRRLRKEDEEED-----FLDD--------S
                       *:.  .::*: :: : . *::  ..: ***     ** *

KAG1050648 TAKEKKVLVPLPLRIILCNL------FADARFFCYTTHRPGWL----TCAIN--------
Homo_sapie TVNPSKCGCPFALKMAACQLLLEITTFLRETFSCLPRPRTEPLVDLESCRLRLDPELDRH
           *.: .*   *:.*::  *:*      *    * * .  *.  *    :* :.

KAG1050648 -------------------------------------------W-ATQPRTNDIG-SQAI
Homo_sapie RYERKISFAGVLDENEDSKDSLHSSSHTLKSDAGVEEKKEGSPWSASEPSIEPEGMSNAG
                                                      * *::*  :  * *:*

KAG1050648 SETFLDENLS-----------ETGFCSN-DMSLPVLNANHIEEALIMFQRMQNTYLDVVE
Homo_sapie AEENYHRNMSWLHVMILLCNQQSFICTHVDYCHPHCYLHHSRSCARLVRAIKLLYGDSVD
           :*   ..*:*           :: :*:: * . *    :* ...  :.: ::  * * *:

KAG1050648 YFRDESSTSHGDSHGTRQAASSLLFSNTQMTENLKYGRRHLAETMYPINQSSSAALNLNP
Homo_sapie SLRESSNISSVALRGKKQKECS--DKSCLRTPSLK-------------KRVSDANL----
            :*:.*. *    :*.:*  .*   ..   * .**             :: *.* *

KAG1050648 PEFDHLENNEGESNSIMLDLAKHKFDGIEEINQDPFGSVFSLLAAVYSTLSSHELGRLLK
Homo_sapie ---------EGKKDSGML-----KYIRLQVMSLSP--APLSLLIKAAPILTEEMYGDIQP
                    **:.:* **     *:  :: :. .*  : :***  . . *:..  * :

KAG1050648 PLWKTYMD-DKKAASFTPAAFLLMECGEKIPKLTTEMF------SQALQRLNILNQSYIQ
Homo_sapie AAWELLLSMDEHMAGAAAAMFLL--CAVKVPEAVSDMLMSEFHHPETVQRLNAVLKFHTL
           . *:  :. *:: *. :.* ***  *. *:*: .::*:      .:::**** : : :

KAG1050648 ISSRNKPF-RGDGGA---FATP------FLPTD-LGSNEFSLDEPRWMSKLKNTSNLPIE
Homo_sapie WRFRYQVWPRMEEGAQQIFKIPPPSINFTLPSPVLGMPSVPMFDPPWVPQCSGSVQDPI-
              * : : * : **   *  *       **:  **  ...: :* *:.: ..: : **

KAG1050648 LKRQIQELGWNEDDN--------------AE------EQEAFKKAL--------------
Homo_sapie ----------NEDQSKSFSARAVSRSHQRAEHILKNLQQEEEKKRLGREASLITAIPITQ
                     ***:.              **      :**  ** *

KAG1050648 --------TP------------------LGLLPSLFLKDDNEDTNDADNPNIVVIRERGK
Homo_sapie EACYEPTCTPNSEPEEEVEEVTNLASRRLSVSPSC--------TSSTSHRNYSF--RRGS
                   **                  *.: **         *..:.: *  .  .**.

KAG1050648 IVDI-SKIITRRKRTATVHS----------------FTAMFLTTVDLLMD-----DFAGV
Homo_sapie VWSVRSAVSAEDEEHTTEHTPNHHVPQPPQAVFPACICAAVLPIVHLMEDGEVREDGVAV
           : .: * : :. :. :* *:                : * .*. *.*: *     * ..*

KAG1050648 NNVLRELIEQFLKDDPALFLRAFLNNLG-KHKADSLKDTLTRIRYIISMQRKL-------
Homo_sapie SAVAQQVLWNCLIEDPSTVLRHFLEKLTISNRQDEL----------MYMLRKLLLNIGDF
           . * :::: : * :**: .** **::*  .:: *.*          : * ***

KAG1050648 PPGLTHILFNYLAGMLKWIVRENQPDGLKFMTLVHPILAELTLSTNDLSVRDL----RKN
Homo_sapie PAQTSHILFNYLVGLIMYFVRTPCEWGMDAISATLTFLWEVVGYVEGLFFKDLKQTMKKE
           *.  :*******.*:: ::**     *:. :: . .:* *:.  .:.* .:**    :*:

KAG1050648 KIE-HLLVSTG----KFWFTHEQPADMFPRYLS-NEQHSFAVL--------NIPQQI---
Homo_sapie QCEVKLLVTASMPGTKTLVVHGQNECDIPTQLPVHEDTQFEALLKECLEFFNIPESQSTH
           : * :***::.    *  ..* *    :*  *. :*: .* .*        ***:.

KAG1050648 FFVACLRISHIQFLINYLARFPREVYAVKKTLQDYEPM-------PVPGSKWETRQLMEK
Homo_sapie YFLMDKRWN----LIHYNKTYVRDIYPFRRSVS---PQLNLVHMHPEKG-----QELIQK
           :*:   * .    **:*   : *::*..::::.   *        *  *     ::*::*

KAG1050648 AYYPD-----------LSRRKNRQSNHKQ------------------EPLDTE-----DS
Homo_sapie QVFTRKLEEVGRVLFLISLTQKIPTAHKQSHVSMLQEDLLRLPSFPRSAIDAEFSLFSDP
             :.            :*  ::  : ***                  ..:*:*     *.

KAG1050648 QQHFDKEMLSL--MRARIWLRFIDVLLNGLNKNYNDHDELERILQGVNTIMIEHSHDFGI
Homo_sapie QA--GKELFGLDTLQKSLWIQLLEEMFLGMPSEFPWGDEIMLFLNVFNGALILHPEDSAL
           *   .**::.*  ::  :*::::: :: *: .::   **:  :*: .*  :* *..* .:

KAG1050648 IGQILILYTRMVTKFKRLFKSNRGYIIFLPALFKVFCESEDNPQITSAITFAWCRFYAVH
Homo_sapie LRQYAATVINTAVHFNHLF-SLSGYQWILPTMLQVYSDYESNPQLRQAIEFACHQFYILH
           : *      . ..:*::** *  **  :**::::*:.: *.***: .** **  :** :*

KAG1050648 EEAFVFQMLGALVPNILMIYGQSDEVGHWMIENLFKLMQAMDNPPHLGSAPDTLGLQLQV
Homo_sapie RKPFVLQLFASVAP-------------------LLEFPDAANNGPSKGVSAQCLFDLLQ-
           .:.**:*::.::.*                   *::: :* :* *  * :.: *   **

KAG1050648 ELDDHERSIQDRIDTASSQVSSALSVSIFKPLARGVTAPISQMEISTYNNR--PFKLEDF
Homo_sapie SLEGETTDILDILELVKAE------------------KPLKSLDF-CYGNEDLTFSISEA
           .*:..  .* * :: ..::                   *:..:::  *.*.  .*.:.:

KAG1050648 IKLFLTVIAYDPGSLRAEQFVKVFTFLIPFFLEQSHLKSMMHE----------GASALVN
Homo_sapie IKLCVTVVAYAPESFRSLQMLMVLEALVPCYLQKLKRQTSQVETVPAAREEIAATAALAT
           *** :**:** * *:*: *:: *:  *:* :*:: : ::   *          .::**..

KAG1050648 VFVKFSKSVKSV------------------VTTNSHTTSGEYQGISSSDSRLPGSNNAAN
Homo_sapie SLQALLYSVEVLTRPMTAPQMSRCDQGHKGTTTANHTMS-----------------SGVN
            :  :  **: :                  .** .** *                 ...*

KAG1050648 TGPKAESA------------QNAYGKQWQQNDRFTIKQEF-------ITLVHKYMKHGGT
Homo_sapie TRYQEQGAKLHFIRENLHLLEEGQGIPREELDERIAREEFRRPRESLLNICTEFYKHCGP
           *  : :.*            ::. *   :: *.   ::**       :.:  :: ** *.

KAG1050648 -------------------LSEVSHEKMAQIIRIVLR----DYRNIKGLTVK--------
Homo_sapie RLKILQNLAGEPRVIALELLDVKSHMRLAEIAHSLLKLAPYDTQTMESRGLRRYIMEMLP
                              *.  ** ::*:* : :*:    * :.::.  ::

KAG1050648 -TDWIKDYLVVSMYSMVDTRNYTKSLKKVLSEIYA----QYRVQWKTVDASDLYEGLAIM
Homo_sapie ITDWTAEAVRPALILIL------KRLDRMFNKIHKMPTLRRQVEWEP--ASNLIEGVCLT
            ***  : :  ::  ::      * *.:::.:*:     : :*:*:.  **:* **:.:

KAG1050648 LEQGQGKAIM----------MNDLAGLVKEKFVSFGLTMATLTDWQNNTKGQVKFCNSLV
Homo_sapie LQRQPIISFLPHLRSLINVCVNLVMGVVGPSSVADGLPLLHLSPYLSPP---LPFSTAVV
           *::    :::          :* : *:*  . *: **.:  *: : . .   : *..::*

KAG1050648 RLIIAILENSSQD-----VFAEIERQVPSVGLIGRVVIPLCLQYNLQWEYSTVGISSTSE
Homo_sapie RLVALQIQALKEDFPLSHVISPFTNQERREGMLLNLLIPFVL---------TVGSGSKD-
           **:   ::  .:*     *:: : .*    *:: .::**: *         *** .*..

KAG1050648 LNSANNWTR------LLNFIAGVCSHTSLLK-SKSSGFSLSQFTTAMGQTNIEDEVDMSD
Homo_sapie ----SPWLEQPEVQLLLQTVINVLLPPRIISTSRSKNFMLES-SPAHCSTPGDAGKDLR-
               . * .      **: : .*   . ::. *:*..* *.. :.*  .*  :   *:

KAG1050648 FPEAKDGKQTPQSVAVLFSLSLIAIKIILIRSHNTFNDLKGS-WSQIAFFIRNTLVFGQT
Homo_sapie ----REGLAESTSQA-----AYLALKVILV----CFERQLGSQWYWLSL---------QV
               ::*   . * *     : :*:*:**:     *:   ** *  :::         *.

KAG1050648 LKLLKAKVGRSDSYNLASPSDVLSPGMPSPSFFSPLGLSSPGASPYLSSNNPSLSPDLRA
Homo_sapie KEMALRKVG---------------------------GLA---------------------
            ::   ***                           **:

KAG1050648 NFSSNAPLGTGTIYDFTTWRFLEFVVCFRTPLFVLLKDFI-----------HEKL-----
Homo_sapie -----------------LWDFLDFIVRTRIPIFVLLRPFIQCKLLAQPAENHEELSARQH
                             * **:*:*  * *:****: **           **:*

KAG1050648 ----------------------------------------------GQMGPNVGSYRNSL
Homo_sapie IADQLERRFIPRPLCKSSLIAEFNSELKILKEAVHSGSAYQGKTSISTVGTSTSAYRLSL
                                                         . :*....:** **

KAG1050648 YS---------------------------PRASFNFPQNENTRWKSWGAEHSSVNTSGDG
Homo_sapie ATMSRSNTGTGTVWEQDSEPSQQASQDTLSRTDEEDEENDSISMPSVVSEQEAYLLSAIG
            :                           .*:. :  :*:.    *  :*:.:   *. *

KAG1050648 QSNANASAHSIVIPKLNIEDTGGVDAETLPTFGLGLHKSNSSASPGHVSHSPTGYSETDS
Homo_sapie RRRFSSHVSSMSVPQAEV---GMLPSQSEPNV---LDDSQGLAAEGSLSRVASIQSEPGQ
           : . .: . *: :*: ::   * : ::: *..   *..*:. *: * :*: .:  **...

KAG1050648 S----------------------SPRSPTTNPFYKQQQHMS-----------PGADVSKE
Homo_sapie QNLLVQQPLGRKRGLRQLRRPLLSRQKTQTEPRNRQGARLSTTRRSIQPKTKPSADQKRS
           .                      * :.. *:*  :*  ::*           *.** .:.

KAG1050648 NNLHLLQAETITSVVNVQVAVGF----------------KPAL---PWITMD-----SRD
Homo_sapie VTFIEAQPEPAAAPTDALPATGQLQGCSPAPSRKPEAMDEPVLTSSPAIVVADLHSVSPK
            .:   *.*. :: .:.  *.*                 :*.*   * *.:      * .

KAG1050648 RSKPWNKKEAVARIANEWQ-------LLLQLST-------ET-------DPLVNKANNNG
Homo_sapie QSENFPTEEGEKEEDTEAQGATAHSPLSAQLSDPDDFTGLETSSLLQHGDTVLHISEENG
           :*: : .:*.  .  .* *       *  ***        **       *.::: :::**

KAG1050648 S-----SANISASPL-------------V
Homo_sapie MENPLLSSQFTFTPTELGKTDAVLDESHV
                 *:::: :*              *


UNC79
>XP_012053306.1 hypothetical protein CNAG_06362 [Cryptococcus neoformans var. grubii H99]
MLSVPEGNTNDDARKALGIRPGGQPPPQLKLDHTNFEDIPRSPALDYPPSPRSPTSSKLPSSPIPPSPTSPEPSRTFRRATRVGRHVLSTKKIQKISSSRGLYETQKLLLYLLDTLEQRETAPDILDRAAISAREISGRLKGKGKGKVLRLGHAIAAAASSSSSIGSSRTHHAMTSTTGVGSEDYEDHIVLDEGDWDTEASYNLVEQTRGLFVLAEKQQLDLFADTGDNDVPILDATPVKSKRRAGRFSSVSPSVAFTKGSSQAAPSSPDISFASTLSPSRTSTSTIPSIPGPYLLERTLNVLQSLLSVDCLHRTRTFRPLCPPNALQAACLDIAAYLYQKGEVGTKVRVVGMVIDGFYGMGEGMEERICEWLEGRMGDLLGRLARERGDFKKPKSDDVEWIDPFSKQDSNTSRNAALPTFAISSESSDSVLRKVSGTQGWMRFSPTSPSFSMFHHGILGLLSIQSLRDASSTAVEIAALVPRILMAITSTIDLTQSKLTTIYRVHRLLSLVLNAKPDSSLDLLTIIAHAPPLSRRTAAEILATFYPNAAGHNTVARRLPSSSYVAQRTKWETGQFGVLGEDETEEHHFVPWRVSSHDDPTAEVVRCEVCESEIHGFGIKCTMCKGQRHLRCYDGSGKTRKGVYQYDVVTLSPTSTSSQLSHAKFCPSLPRLEEVLLSPPHGSECTHRKAGQHRLDLIHLFTLTLCDECHFPLWGVSKQGYTCQNGCQRFFHSHCLEKMEETGVGECRYGREVVVDEISEAGNNPFVITRDKLQSSFHQHYNGFLPSKDELKGKTFDEIGVIFGTIWIQYQLIQNGLSSGSLRVSNEDKKATSDPLGLKPILKTYEELLQSHEQAISTAAADFTHAATLEKPLGTGYLFSEKFLGYCTALIRAPTTSSSTLSREGEHLGDGFLTPQGLSPSLEEGDKVDRCYETLPLGIIVQSLATDLSLSNKVAATAFLNHLSLVGFISIFDTASLTSQYLAKRREIKVSFTLPLLMDSSPNTELLVLAIEALLDDLDLTMNEQGLRLLVTRAWPSLLCAPYALERLGKACVSWVISEEDCLRDIIKKYASKHRRIPGVRPAAGAQKGSVDMYKQDRQRLRRMFVEPWLKALHDQDPALYVHIVYEQCKLVAANVAMEDLSGASEEQVASRMAGVAVSKMTSMGEAGMLFSTVMELLTAWLEDLGPLADHDVAYRALPRLLNHQPADSADIFGLSLSTAQAGPADLARVCRWMRVLSYSGVEIPWELLLSLVDLQAGSPAFTFDGSVGGASAEAKLDLVIAINSNRAVIEPQTFASACSRLAVGVFCDMGKHEKMEPLELELVKRTMLLILQAYGVPVDEVAETTLGVGMPSTGQPPTITKKRQSPIVNVRFPLNADMVVGAATLLERTSCPDEMVLDFLWLLSTKAGMVDDPVGFLHHTCSKLYEMIWPLIGLPIDRRSRARVLLKLLSVNSAPLERIIHAQLEFSPEARAQARERLLIFILELADTSVNYELSNWRAAMVGLVLLFFDVLLDPRDVTPDNIIILKTLQPTQLNAMSMCFEEHLVKSSDERRLVLLSRLSRLRMNVPQWAIISWTTIDELLAEEVASLTQFKRADQSQTDSGRVDSQNVAYSLIFLGLEMLAAGVPITWIAAQRFQQRVAAACALPWFNPPVFTAVVLPGLRSVLDSPIRIMISGETFESKVKKTVLVGSLFVPVVIDFAQELKKYDVAVQRILLDILMVTFFKQDVTRVELSTYSAVQKVADFVLTGECSENRLLALQILQIAVTKVERDKIIRAVPPAFNTVAKVLVKELEAEYGDPAVVEQSQLFLRNIVKGFGRSGLYLQLFRNESESYDLSSPHETSLAKALQILHNEQEREALPQASLFDNVFHDLLDVTKRPRQQVIHIIEAFARFATSFEGHLSEEAAQDFGTFVNRLSKLIAEWSFDFDPNPILQSCAKILDRTPSASLVPLLSQVSTILNHCMGHFAVKRATAIRLLESGETASRKAQTENQIRIVLFEMAGAAINGLPVTPEGLYTLLKYLTSDAFSRPAPDSRVSSEQIRVLSDASPGCVHILLNGHPALESNLMSAEVLVAVLNEAGTVLCQTEMVVPGAISRNVTQLTLDAASSQVNFFLYLLLSSLDLTMGPARSRLLSLYPLLSRATSLCLRAASDYMSLQDVEGHGTFLISLAFMVMRLALLAVRDGPVSNEDREKGAGDEDAVDILWTRIWPEWIRLFRLSFDPNCVNGALRTVTHSVFLDLLTFLGGIYSPILLRHTESLSVNLAMLVQYQETLGVAPSGKLQKATQIMEKVGTHSVGGVADRMGMMESLKADLLAMERMKILNK


>Homo_sapiens_NP_001333147.1
MSTKAEQFASKIRYLQEYHNRVLHNIYPVPSGTDIANTLKYFSQTLLSILSRTGKKENQDASNLTVPMTMCLFPVPFPLTPSLRPQVSSINPTVTRSLLYSVLRDAPSERGPQSRDAQLSDYPSLDYQGLYVTLVTLLDLVPLLQHGQHDLGQSIFYTTTCLLPFLNDDILSTLPYTMISTLATFPPFLHKDIIEYLSTSFLPMAILGSSRREGVPAHVNLSASSMLMIAMQYTSNPVYHCQLLECLMKYKQEVWKDLLYVIAYGPSQVKPPAVQMLFHYWPNLKPPGAISEYRGLQYTAWNPIHCQHIECHNAINKPAVKMCIDPSLSVALGDKPPPLYLCEECSERIAGDHSEWLIDVLLPQAEISAICQKKNCSSHVRRAVVTCFSAGCCGRHGNRPVRYCKRCHSNHHSNEVGAAAETHLYQTSPPPINTRECGAEELVCAVEAVISLLKEAEFHAEQREHELNRRRQLGLSSSHHSLDNADFDNKDDDKHDQRLLSQFGIWFLVSLCTPSENTPTESLARLVAMVFQWFHSTAYMMDDEVGSLVEKLKPQFVTKWLKTVCDVRFDVMVMCLLPKPMEFARVGGYWDKSCSTVTQLKEGLNRILCLIPYNVINQSVWECIMPEWLEAIRTEVPDNQLKEFREVLSKMFDIELCPLPFSMEEMFGFISCRFTGYPSSVQEQALLWLHVLSELDIMVPLQLLISMFSDGVNSVKELANQRKSRVSELAGNLASRRVSVASDPGRRVQHNMLSPFHSPFQSPFRSPLRSPFRSPFKNFGHPGGRTIDFDCEDDEMNLNCFILMFDLLLKQMELQDDGITMGLEHSLSKDIISIINNVFQAPWGGSHTCQKDEKAIECNLCQSSILCYQLACELLERLAPKEESRLVEPTDSLEDSLLSSRPEFIIGPEGEEEENPASKHGENPGNCTEPVEHAAVKNDTERKFCYQQLPVTLRLIYTIFQEMAKFEEPDILFNMLNCLKILCLHGECLYIARKDHPQFLAYIQDHMLIASLWRVVKSEFSQLSSLAVPLLLHALSLPHGADIFWTIINGNFNSKDWKMRFEAVEKVAVICRFLDIHSVTKNHLLKYSLAHAFCCFLTAVEDVNPAVATRAGLLLDTIKRPALQGLCLCLDFQFDTVVKDRPTILSKLLLLHFLKQDIPALSWEFFVNRFETLSLEAQLHLDCNKEFPFPTTITAVRTNVANLSDAALWKIKRARFARNRQKSVRSLRDSVKGPVESKRALSLPETLTSKIRQQSPENDNTIKDLLPEDAGIDHQTVHQLITVLMKFMAKDESSAESDISSAKAFNTVKRHLYVLLGYDQQEGCFMIAPQKMRLSTCFNAFIAGIAQVMDYNINLGKHLLPLVVQVLKYCSCPQLRHYFQQPPRCSLWSLKPHIRQMWLKALLVILYKYPYRDCDISKILLHLIHITVNTLNAQYHSCKPHATAGPLYSDNSNISRYSEKEKEEDSVFDESDIHDTPTGPCNKESQTFFARLKRIGGSKMVKYQPVEMNVQRSEIELAEYRETGALQDSLLHCVREESIPKKKLRSFKQKSLDIGNADSLLFTLDEHRRKSCIDRCDIEKPPTQAAYIAQRPNDPGRSRQNSATRPDNSEIPENPAMEGFPDARRPVIPEVRLNCMETFEVKVDSPVKPAPKEDLDLIDLSSDSTSGPEKHSILSTSDSDSLVFEPLPPLRIVESDEEEETMNQGDDGPSGKNAASSPSVPSHPSVLSLSTAPLVQVSVEDCSKDFSSKDSGNNQSAGNTDSALITLEDPMDAEGSSKPEELPEFSCGSPLTLKQKRDLLQKSFALPEMSLDDHPDPGTEGEKPGELMPSSGAKTVLLKVPEDAENPTESEKPDTSAESDTEQNPERKVEEDGAEESEFKIQIVPRQRKQRKIAVSAIQREYLDISFNILDKLGEQKDPDPSTKGLSTLEMPRESSSAPTLDAGVPETSSHSSISTQYRQMKRGSLGVLTMSQLMKRQLEHQSSAPHNISNWDTEQIQPGKRQCNVPTCLNPDLEGQPLRMRGATKSSLLSAPSIVSMFVPAPEEFTDEQPTVMTDKCHDCGAILEEYDEETLGLAIVVLSTFIHLSPDLAAPLLLDIMQSVGRLASSTTFSNQAESMMVPGNAAGVAKQFLRCIFHQLAPNGIFPQLFQSTIKDGTFLRTLASSLMDFNELSSIAALSQLLEGLNNKKNLPAGGAMIRCLENIATFMEALPMDSPSSLWTTISNQFQTFFAKLPCVLPLKCSLDSSLRIMICLLKIPSTNATRSLLEPFSKLLSFVIQNAVFTLAYLVELCGLCYRAFTKERDKFYLSRSVVLELLQALKLKSPLPDTNLLLLVQFICADAGTKLAESTILSKQMIASVPGCGTAAMECVRQYINEVLDFMADMHTLTKLKSHMKTCSQPLHEDTFGGHLKVGLAQIAAMDISRGNHRDNKAVIRYLPWLYHPPSAMQQGPKEFIECVSHIRLLSWLLLGSLTHNAVCPNASSPCLPIPLDAGSHVADHLIVILIGFPEQSKTSVLHMCSLFHAFIFAQLWTVYCEQSAVATNLQNQNEFSFTAILTALEFWSRVTPSILQLMAHNKVMVEMVCLHVISLMEALQECNSTIFVKLIPMWLPMIQSNIKHLSAGLQLRLQAIQNHVNHHSLRTLPGSGQSSAGLAALRKWLQCTQFKMAQVEIQSSEAASQFYPL


CLUSTAL format alignment by MAFFT (v7.511)


XP_0120533 MLS-------------------------VPEG----NT----------------------
Homo_sapie MSTKAEQFASKIRYLQEYHNRVLHNIYPVPSGTDIANTLKYFSQTLLSILSRTGKKENQD
           * :                         **.*    **

XP_0120533 ------------------------------------------------------------
Homo_sapie ASNLTVPMTMCLFPVPFPLTPSLRPQVSSINPTVTRSLLYSVLRDAPSERGPQSRDAQLS


XP_0120533 ----------------------------------------------NDD-----------
Homo_sapie DYPSLDYQGLYVTLVTLLDLVPLLQHGQHDLGQSIFYTTTCLLPFLNDDILSTLPYTMIS
                                                         ***

XP_0120533 -----------------------------ARK----------------------------
Homo_sapie TLATFPPFLHKDIIEYLSTSFLPMAILGSSRREGVPAHVNLSASSMLMIAMQYTSNPVYH
                                        :*:

XP_0120533 ------------------------------------------------------------
Homo_sapie CQLLECLMKYKQEVWKDLLYVIAYGPSQVKPPAVQMLFHYWPNLKPPGAISEYRGLQYTA


XP_0120533 --------------------------ALGIRPGGQPPP---------QLKLDHTNFEDIP
Homo_sapie WNPIHCQHIECHNAINKPAVKMCIDPSLSVALGDKPPPLYLCEECSERIAGDHSEWLIDV
                                     :*.:  *.:***         ::  **:::

XP_0120533 RSPALDYPPSPRSPTSSKLPSSPIPPSPTSPEPSRTFRRATRVGRHVLSTKKIQKI--SS
Homo_sapie LLPQAEISAICQKKNCSSHVRRAVVTCFSAGCCGRHGNRPVRYCKRCHSNHHSNEVGAAA
             *  : ..  :. ..*.    .: .. ::   .*  .*..*  ::  *.:: :::  ::

XP_0120533 SRGLYETQKLLLYLLDTLEQRETAPDILDRAAISAREISGRLKGKGKGKVLRLGHAIAAA
Homo_sapie ETHLYQTSP------PPINTRECGAEEL----VCAVEAVISLLKEAEFHAEQREHEL---
           .  **:*.        .:: ** ..: *    :.* *    *  :.: :. :  * :

XP_0120533 ASSSSSIGSSRTHHAMTSTTGVGSED--YEDHIVLDEGDW--------------------
Homo_sapie -NRRRQLGLSSSHHSLDNADFDNKDDDKHDQRLLSQFGIWFLVSLCTPSENTPTESLARL
            .   .:* * :**:: .:   ..:*  :::::: : * *

XP_0120533 ---------------DTEASYNLVEQTRGLFVLAEKQQLDLFADTG-DNDVPILDATPVK
Homo_sapie VAMVFQWFHSTAYMMDDEVG-SLVEKLKPQFV---TKWLKTVCDVRFDVMVMCLLPKPME
                          * *.. .***: :  **   .: *. ..*.  *  *  * ..*::

XP_0120533 SKRRAGRFSSVSPSVAFTKGSSQAAPSSPDISFASTLSPSRTSTSTIPSIPGPYLLERTL
Homo_sapie FARVGGYWD---------KSCSTVTQLKEGLNRILCLIPYNVINQSVWECIMPEWLEAIR
             * .* :.         *..* .:  . .:.    * * .. ..:: .   *  **

XP_0120533 NVLQSLLSVDCLHRTRTFR---------PLCP-------------------PNALQAACL
Homo_sapie TEVPD-------NQLKEFREVLSKMFDIELCPLPFSMEEMFGFISCRFTGYPSSVQEQAL
           . : .       :: : **          ***                   *.::*  .*

XP_0120533 DIAAYLYQKGEVGTKVRVVGMVIDGFYGMGEGMEERICEWLEGRMGDLLGRLARERGDFK
Homo_sapie LWLHVLSELDIMVPLQLLISMFSDGVNSVKELANQR-----KSRVSELAGNLASRR----
                * : . : .   ::.*. **. .: *  ::*     :.*:.:* *.** .*

XP_0120533 KPKSDDVEWIDPFSKQDSNTSRNAALPTFAISSESSDSVLRKVSGTQGWMRFSPTSPSFS
Homo_sapie ---------VSVASDPGRRVQHNMLSPFHSPFQSPFRSPLR-----------SPFRSPFK
                    :.  *. . ...:*   * .:  ...  * **           **  ..*.

XP_0120533 MFHHGILGLLSIQSLRDASSTAVEIAALVPRILM--------AITSTIDLTQSKLTTIYR
Homo_sapie NFGHP--GGRTIDFDCEDDEMNLNCFILMFDLLLKQMELQDDGITMGLEHSLSK-DIISI
            * *   *  :*:   : ..  ::   *:  :*:        .**  :: : **   *

XP_0120533 VHRLLSLVLNAKPDSSLDLLTI---IAHAPPLSRRTAAEILATFYPNAAGHNTVARRLPS
Homo_sapie INNVFQAPWGGSHTCQKDEKAIECNLCQSSILCYQLACELLERLAPKEESRLVEPTDSLE
           ::.::.   ...  .. *  :*   :.::. *. : *.*:*  : *:  .: . .    .

XP_0120533 SSYVAQRTKWETGQFGVLGEDETEEHHFVPWRVSSHDDPTAEVVRCEVCESEIHGFGIKC
Homo_sapie DSLLSSRPEFIIGPEGEEEENPASKHGENPGN----------------CTEPVEHAAVK-
           .* ::.*.::  *  *   *: :.:*   * .                * . :.  .:*

XP_0120533 TMCKGQRHLRCYDGSGKTRKGVYQYDVVTLSPTSTSSQLSHAKFCPS---LPRLEEVLLS
Homo_sapie --------------NDTERKFCYQQLPVTLRLIYTIFQ-EMAKFEEPDILFNMLNCLKIL
                         ... **  **   ***    *  * . ***  .   :  *: : :

XP_0120533 PPHGSECTHRKAGQHRLDLIHLFTLTLCDECHFPLWGVSKQGYTCQNGCQRFFHSHCLEK
Homo_sapie CLHG-ECLYIARKDHPQFLAYIQDHMLIAS----LWRVVKSEFSQLSSLAVPLLLHALSL
             ** ** :    :*   * ::    *  .    ** * *. ::  ..    :  *.*.

XP_0120533 MEETGV-------------GECRYGREVVVDEISEAGNNPFVITRDKLQSSFHQHYNGFL
Homo_sapie PHGADIFWTIINGNFNSKDWKMRFEAVEKVAVICRFLDIHSVTKNHLLKYSLAHAFCCFL
            . :.:              : *:     *  *..  :   * ... *: *: : :  **

XP_0120533 PSKDELKGKTFDEIGVIFGTIWIQYQLIQNGLSSGSLRVSNEDKKATSDPLGLKPILKTY
Homo_sapie TAVEDVNPAVATRAGLLLDTI------KRPALQGLCLCLDFQFDTVVKD----RPTILSK
           .: ::::  .  . *:::.**       : .*.. .* :. : .....*    :* : :

XP_0120533 EELLQSHEQAISTAAADF----THAATLEKPLGTGYLFSEKFLGYCTALIRAPTTSSSTL
Homo_sapie LLLLHFLKQDIPALSWEFFVNRFETLSLEAQL---HLDCNKEFPFPTT-ITAVRTNVANL
             **:  :* *.: : :*     .: :**  *   :* .:* : : *: * *  *. :.*

XP_0120533 S-------REGEHLGDGFLTPQGLSPSLE---EGDKVDRCYETLPLGIIVQSLATDLSLS
Homo_sapie SDAALWKIKRARFARNRQKSVRSLRDSVKGPVESKRALSLPETLTSKIRQQSPENDNTIK
           *       :....  :   : :.*  *::   *..:.    ***.  *  **  .* ::.

XP_0120533 NKVAATAFLNHLSLVGFISIF------DTASLTSQYLAKRREIKVSFTLPLLMDSSPNTE
Homo_sapie DLLPEDAGIDHQTVHQLITVLMKFMAKDESSAESDISSAKAFNTVKRHLYVLLGYDQQEG
           : :.  * ::* ::  :*:::      * :*  *:  : :   .*.  * :*:. . :

XP_0120533 LLVLA-------------IEALLDDLDLTMNEQGLRLLVTRAWPSLLCAPYALERLGKAC
Homo_sapie CFMIAPQKMRLSTCFNAFIAGIAQVMDYNIN------------------------LGKHL
            :::*             * .: : :* .:*                        ***

XP_0120533 VSWVISEEDCLRDIIKKYASKHRRIPGVRPAAGAQKGSVDMYKQDRQRLRRMFVEP----
Homo_sapie LPLVVQ--------VLKYCS----CPQLR-----------HYFQQPPRCSLWSLKPHIRQ
           :. *:.        : **.*     * :*            * *:  *     ::*

XP_0120533 -WLKAL---------HDQD-PALYVHIV----------YEQCK-------LVAANVAMED
Homo_sapie MWLKALLVILYKYPYRDCDISKILLHLIHITVNTLNAQYHSCKPHATAGPLYSDNSNISR
            *****         :* * . : :*::          *..**       * : *  :.

XP_0120533 LSGASEEQVA-----------------------SRMAGVAVSKMTSMGEAGMLFSTVMEL
Homo_sapie YSEKEKEEDSVFDESDIHDTPTGPCNKESQTFFARLKRIGGSKMVKYQPVEMNVQR-SEI
            *  .:*: :                       :*:  :. ***..   . * ..   *:

XP_0120533 LTAWLEDLGPLAD---HDVAYRALPR----------------------LLNHQPADSADI
Homo_sapie ELAEYRETGALQDSLLHCVREESIPKKKLRSFKQKSLDIGNADSLLFTLDEHRRKSCIDR
             *  .: *.* *   * *  .::*:                      * :*:  .. *

XP_0120533 FGLSLSTAQAG-----PADLARVCRWMRVLSYSGVEIPWELLLSLVDLQAGSPAF-----
Homo_sapie CDIEKPPTQAAYIAQRPNDPGR-SRQNSATRPDNSEIPENPAMEGFP-DARRPVIPEVRL
            .:. ..:**.     * * .* .*   .   .. *** :  :. .  :*  *.:

XP_0120533 ----TFDGSVGGASAEA---KLDLVIAINSNRAVIEPQTFASACSRLAVGVFCDMGKHEK
Homo_sapie NCMETFEVKVDSPVKPAPKEDLDLIDLSSDSTSGPEKHSILSTSDS-------DSLVFEP
               **: .*...   *   .***:   ... :  * ::: *:..        *   .*

XP_0120533 MEPLELELVKRTMLLILQAYGVPVDEVAETTLGVGMPSTGQPPTITKKRQSPIVNVRFP-
Homo_sapie LPPLRIVESDEEEETMNQGDDGPSGKNAASS-----PSVPSHPSVLSLSTAPLVQVSVED
           : **.:   ..    : *. . * .: * ::     **. . *:: .   :*:*:* .

XP_0120533 LNADMVVGAATLLERTSCPDEMVLDFLWLLSTKAGMVDDPVGFLHHTCSKLYEMIWPLIG
Homo_sapie CSKDFSSKDSGNNQSAGNTDSALI-----------TLEDPMD--AEGSSKPEELPEFSCG
            . *:    :   : :. .*. ::            ::**:.   . .**  *:     *

XP_0120533 LPIDRRSRARVLLKLLSVNSAPLERIIHAQLEFSPEAR---AQARERLLIFILELADTSV
Homo_sapie SPLTLKQKRDLLQKSFALPEMSLDDHPDPGTEGEKPGELMPSSGAKTVLLKVPEDAENPT
            *:  :.:  :* * ::: . .*:   ..  * .  ..   :.. : :*: : * *:...

XP_0120533 NYELSNWRAAMVGLVLLFFDVLLDP-RDVTPDNIIILKTLQPTQLNAMSMCFEEHLVKSS
Homo_sapie ESEKPDTSAES--------DTEQNPERKVEED-------------GAEESEFKIQIVPRQ
           : * .:  *          *.  :* *.*  *             .* .  *: ::*  .

XP_0120533 DERRLVLLSRLSRLRMNVPQWAIISWTTIDELLAEE--------VASLTQFKRADQSQTD
Homo_sapie RKQRKIAVSAIQR------EYLDISFNILDKLGEQKDPDPSTKGLSTLEMPRESSSAPTL
            ::* : :* :.*      ::  **:. :*:*  ::        :::*   :.:..: *

XP_0120533 SGRVDSQNVAYSLIFLGLEMLAAGVPITWIAAQRFQQRVAAACALPWFNPPVFTAVVLPG
Homo_sapie DAGV-PETSSHSSISTQYRQMKRGSLGVLTMSQLMKRQLEHQSSAPHNISNWDTEQIQPG
           .. * .:. ::* *    . :  *   .   :* :::::   .: *   .   *  : **

XP_0120533 LRS----------VLDSPIRIMISGETFESKVKKTVLVGSLFVPVVIDFAQELKKYDVAV
Homo_sapie KRQCNVPTCLNPDLEGQPLR--MRGATKSSLLSAPSIV-SMFVPAPEEFTDEQPT-----
            *.          : ..*:*  : * * .* :. . :* *:***.  :*::*  .

XP_0120533 QRILLDILMVTFFKQDVTRVELSTYSAVQKVADFVLTGECSENRLLALQILQIAVTKVER
Homo_sapie --------VMTDKCHDCGAILEEYDEETLGLAIVVLSTFIHLSPDLAAPLLLDIMQSVGR
                   ::*   :*   :  .  . .  :* .**:     .  **  :*   : .* *

XP_0120533 DKIIRAVPPAFNTVAKVLVKELEAEYGDPAVVEQSQLFLRNIVKGFGRSGLYLQLFRNES
Homo_sapie ----LASSTTFSNQAESMMVP-----GNAAGVAKQ--FLRCIFHQLAPNGIFPQLFQSTI
                * ..:*.. *: ::       *:.* * :.  *** *.: :. .*:: ***:.

XP_0120533 E---------SYDLSSPHETSLAKALQILHNEQEREALPQASLFDNVFHDLLDVTKRPRQ
Homo_sapie KDGTFLRTLASSLMDFNELSSIAALSQLLEGLNNKKNLPAGGA-----------------
           :         *  :.  . :*:*   *:*.. :::: ** ..

XP_0120533 QVIHIIEAFARFATSFEGHLSEEAAQDFGTFVNRLSKLIAEWSFDFDPNPILQSCAKI--
Homo_sapie ----MIRCLENIATFMEALPMDSPSSLWTTISNQFQTFFAKLPCVLPLKCSLDSSLRIMI
               :*..: .:** :*.   :..:. : *: *::..::*: .  :  :  *:*. :*

XP_0120533 -LDRTPSASLV-PLLSQVSTILNHCMGHFAVKRATAIRLLESGETASRKAQTENQI-RIV
Homo_sapie CLLKIPSTNATRSLLEPFSKLLSFVIQNAVFTLAYLVELCGLCYRAFTKERDKFYLSRSV
            * : **:. . .**. .*.:*.. : : ... *  :.*      *  * : :  : * *

XP_0120533 LFEMAGAAINGLPVTPEGLYTLLKYLTSDAFSRPAPDSRVSSEQIRVL---SDASPGCVH
Homo_sapie VLELLQALKLKSPLPDTNLLLLVQFICADAGTKLAESTILSKQMIASVPGCGTAAMECVR
           ::*:  *     *:.  .*  *:::: :** :: * .: :*.: *  :   . *:  **:

XP_0120533 ILLNGHPALESNLMSAEVLVAVLNEAGTVLCQTEMVVPGAISRNVTQLTLDAAS------
Homo_sapie QYINEVLDFMADMHTLTKLKSHMKTCSQPL--HEDTFGGHLKVGLAQIAAMDISRGNHRD
             :*    : ::: :   * : :: ..  *   * .. * :. .::*::    *

XP_0120533 --SQVNFFLYLLLSSLDLTMGP-------ARSRLLSLYPLLS--------RATSLCLRAA
Homo_sapie NKAVIRYLPWLYHPPSAMQQGPKEFIECVSHIRLLSWLLLGSLTHNAVCPNASSPCLPIP
             : :.:: :*  ..  :  **       :: ****   * *        .*:* **  .

XP_0120533 SDYMS----------LQDVEGHGTFLISL-----AFMVMRLALLAVRDGPVSNEDREKGA
Homo_sapie LDAGSHVADHLIVILIGFPEQSKTSVLHMCSLFHAFIFAQLWTVYCEQSAVATNLQNQNE
            *  *          :   *   * :: :     **:. :*  :  .:..*:.: :::.

XP_0120533 GDEDAVDI---LWTRIWPEWIRLF---RLSFDPNCVN-----GALRTVTHSVFLDLLTFL
Homo_sapie FSFTAILTALEFWSRVTPSILQLMAHNKVMVEMVCLHVISLMEALQECNSTIFVKLIP--
            .  *:     :*:*: *. ::*:   :: .:  *::      **:  . ::*:.*:.

XP_0120533 GGIYSPILLRHTESLSVNLAMLVQY-----------------QETLGVAPSGKLQKATQI
Homo_sapie --MWLPMIQSNIKHLSAGLQLRLQAIQNHVNHHSLRTLPGSGQSSAGLAALRKWLQCTQF
             :: *::  : : **..* : :*                  *.: *:*.  *  :.**:

XP_0120533 -MEKVGTHSVGGVADRMGMMESLKADLLAMERMKILNK
Homo_sapie KMAQVEIQSSEAASQFYPL-------------------
            * :*  :*  ..::   :
```
